# A possible mechanistic insight on how Compromised Hydrolysis of Triacylglycerol 7 (CHT7) restrains the involvement of it’s DNA binding CXC domain from quiescence repression

**DOI:** 10.1101/2023.10.23.563394

**Authors:** Manisha Chauhan, Syeda Amna Arshi, Naveen Narayanan, Haseeb Ul Arfin, Amit Sharma

## Abstract

CHT7 is a regulator of quiescence repression and TAG degradation between the nitrogen deprived and the nitrogen replenished states in *Chlamydomonas reinhardtii*. Initially it was thought that the CHT7’s repression activity is managed by its DNA binding CXC domain which is a tandem repeat of two cysteine rich subdomains. Later, it was found that the CXC (CHT7_CXC) domain is effectively dispensable for CHT7’s activities. Rather, CHT7’s predicted protein binding domains are proposed to be involved in gene regulation activities by binding through other repressors in the cell. Yet, it remains unclear why and how CHT7 manages to refrain its own CXC domain from participating in any transcriptional activities. The question becomes more intriguing, because CXC binding regions are available in promoter regions of some of the misregulated genes in the CHT7 mutant (*cht7*). Through the combination of biophysical experiments and molecular dynamics approaches, we have studied the DNA recognition behavior of CHT7_CXC. The results show that CHT7_CXC domain is highly selective towards DNA sequences and this selectivity is imparted due to the differential binding abilities of the CXC subdomains. Further, to understand if the case is - that CXC looses it’s DNA binding capabilities in the vicinity of other repressor molecules, we carried out CHT7_CXC’s DNA binding stability test by simulating the spatial constraint conditions using the AsLOV2- CXC fusion. Our test results show limited ability of CHT7_CXC to withstand steric forces and provide insights to why and how algal cells may hold back CHT7_CXC’s indulgence in quiescence repression.

**Significance:** Microalgae, under nutrient rich conditions, provide biomass. Whereas, nutrient deprivation leads to accumulation of biofuel feedstock, but cells enter quiescence. Net enhancement in feedstock, therefore relies on the precision of the quiescence regulator. In *Chlamydomonas reinhardtii*, CHT7 is a central regulator of quiescence. Surprisingly, rather than using its own DNA binding domain (DBD) for the regulatory activities, CHT7 recruits external transcriptional regulators using its non DBDs. To ensure smooth functioning, CHT7’s DBD must rapidly switch to inactive form. Modifications in DNA binding profiles of DBDs due to non DBDs are seen in transcription factors of many organisms. The switching mechanism discussed could therefore be a generic approach of timely regulation of individual components of the complex transcriptional machineries.

## Introduction

Lack of nutrients put the cellular organisms under stress. To survive, cells enter the quiescence phase and reduce their energy expenditure (1, 2). Specifically, the *Chlamydomonas reinhardtii* cells manage quiescence by reducing activities related to cellular growth, division, photosynthesis, metabolism and translation, and by inducing autophagy and accumulation of carbon concentrating molecules such as triacylglycerol (TAG) (2–5). Whereas, on nutrient replenishment (NR), these cells exit quiescence, utilize the accumulated TAG as an initial burst of energy and restart its growth and division activities (3, 6). Extensive work by Benning and coauthors have shown that the entry and exit from quiescence is regulated through a protein called Compromised Hydrolysis of Triacylglycerol 7 (CHT7) (3, 7, 8). Efficient entry into the quiescence and maintenance of this state are necessary for a successful exit from the quiescence (9, 10). Hence, it is of utmost importance that CHT7 conduct gene regulation activities with high precision.

CHT7, a nuclear protein, houses four predicted protein domains (PPDs: P1-P4) and a DNA binding CXC domain (CHT7_CXC) (**Fig.S1A**). Wherein, CXC domain contains two subdomains, N terminal (CXC_N) and C terminal (CXC_C) connected by a loop. CHT7 maintains its repression activities using its PPDs and the CXC domain is dispensable for its functionality **(7)**. The CXC homologs are found in various plant and animal species and are highly conserved domains (**Fig.S1B)**. CXC proteins for example, CBBP, CPP1, TSO1, MSL2, TESMIN, EZH2 and LIN54 have role in paramutation and heritable gene expression in maize **(11)**, regulation of the leghemoglobin gene expression in Soybean **(12)**, modulates cytokinesis, meristem organization and cell division in *Arabidopsis thaliana* **(13, 14)**, X chromosome’s dosage compensation in *Drosophila melanogaster* males **(15)**, spermatogenesis in mouse **(16)**, transcriptional repression in humans **(17, 18)** and cell cycle regulatory functions in humans **(19)**, respectively. LIN54 is particularly relevant because of its CXC domain’s high sequence identity (56.6%) with CHT7_CXC. It performs cellular functions by recognizing 5’TTYRAA3’ (where Y is pyrimidine and R is purines) stretches in the DNA minor groove **(20)**. Tyr present in its CXC_N subdomain form H-bonds with A6 and complementary nucleotide A5 (A5* i.e. T) and CXC_C forms H bonds with T2 and complementary nucleotide T1 (T1* i.e. A). Any deviation from T1, T2, A5 and A6, as well as the absence of either CXC_N or CXC_C results in no DNA binding for LIN54 **(20)**.

Considering that CXC homologs possess precise DNA binding mechanisms which are exclusively important for the regulation of specific cellular activities, CHT7_CXC may also regulate transcription activities (3). On the contrary, the suggested mode of regulation by CHT7 is, it undergoes transcriptional activities by forming large complexes by binding to the external transcriptional regulators via its PPDs (7). Whereas, its CXC domain has been found dispensable for any transcriptional activity.

A few characterizations in relation to CXC’s DNA binding are required to understand why CHT7 refrains its CXC from participating in transcriptional activities. Thus, we asked in this study does, 1) *C*.*reinhardtii* genome have promoter regions specifically recognizable by the CXC domain, 2) CHT7_CXC possess sequence-specific binding abilities, 3) CHT7 restrains CXC DNA interactions and 4) considering that the CHT7 forms multiprotein complexes, is CXC DNA interactions sufficiently stable to withstand the steric forces due to molecular crowding within these complexes.

Here we examined these scenarios using the biophysical and computational approaches. Our results favour the DNA binding possibilities for the CHT7_CXC and find molecular crowding within the CHT7 complexes a plausible reason behind the non-engagement of CXC in the transcriptional activities.

## Results

### CHT7_CXC possesses sequence-specific DNA recognition ability

To explore the DNA binding ability of CHT7_CXC, promoters were identified (**SI**) for the set of misregulated genes **(3)** in the cht7 mutant. TTYRAA (the LIN54 binding region) could be located in some of the predicted promoters. Out of these, the 27mer stretch of the CDKA1 promoter, that contained in it TTTGAA region, was used for the protein DNA binding studies. The Fluorescence Polarization Assays (FPA) **(21)** were conducted in which CHT7_CXC was titrated against 6-Carboxyfluorescein (6-FAM) tagged DNA substrate (6-FAM-5’-TATCTGGTG**TTTGAA**TTTCTGGATCTG-3’). The sigmoidal increase in fluorescence anisotropy (**Blue, Fig.1A**) demonstrates protein DNA binding with the K_D_ value of 873.38±133.17nM (**Table1**). Binding was also confirmed through Electrophoretic Mobility Shift Assay (EMSA), in which the appearance of a protein-DNA bound fraction (B_D_ and B_P_ in TTTGAA gel **Fig.1C)** indicates a single binding conformation **(22, 23)** for CHT7_CXC-TTTGAA complex. Further, no binding was observed when the binding site was replaced with 5’-**CCTGCC**-3’ (**Red, Fig.1A and 1C**) or when the DNA sequence contained none of the TTYRAA sites (Lane 7, **Fig.S2**). Also, no binding (**Black, Fig.1A;** Lane 1-6, **Fig.S2**) could be observed when the shorter (13 mer) DNA fragment (5’-GGTG**TTTGAA**TTT-3’) was used. These results indicate that CHT7_CXC binding is highly dependent on the binding sequence and the length of the DNA. Since, CXC binding sites could also be located in the promoter regions of the genes other than the cell cycle genes (**Fig.S3)**, their recognition by the CHT7_CXC was confirmed for two photosynthetic genes LHCA6 and LHCB5 (**Fig.S4)**.

**Table 1:** Protein dissociation constants using FPA.

| **DNA** | **Protein** | **KD (nM)** | **SE(n)** | **R<sup>2</sup>** |
| --- | --- | --- | --- | --- |
| **TTTGAA** | **CXC** | **873.38** | **133.17 (3)** | **0.99** |
|  | **CXC_N** | **2633.33** | **906.2 (3)** | **0.99** |
|  | **CXC_C** | **NB** |  |  |
| **CCTGAA** | **CXC** | **368.31** | **133.42 (3)** | **0.96** |
|  | **CXC_N** | **NB** |  |  |
|  | **CXC_C** | **NB** |  |  |
| **CCTGCC** | **CXC** | **NB** |  |  |
| **TTTGCC** | **CXC** | **NB** |  |  |
|  | **CXC_N** | **NB** |  |  |
|  | **CXC_C** | **NB** |  |  |

**Fig1:**
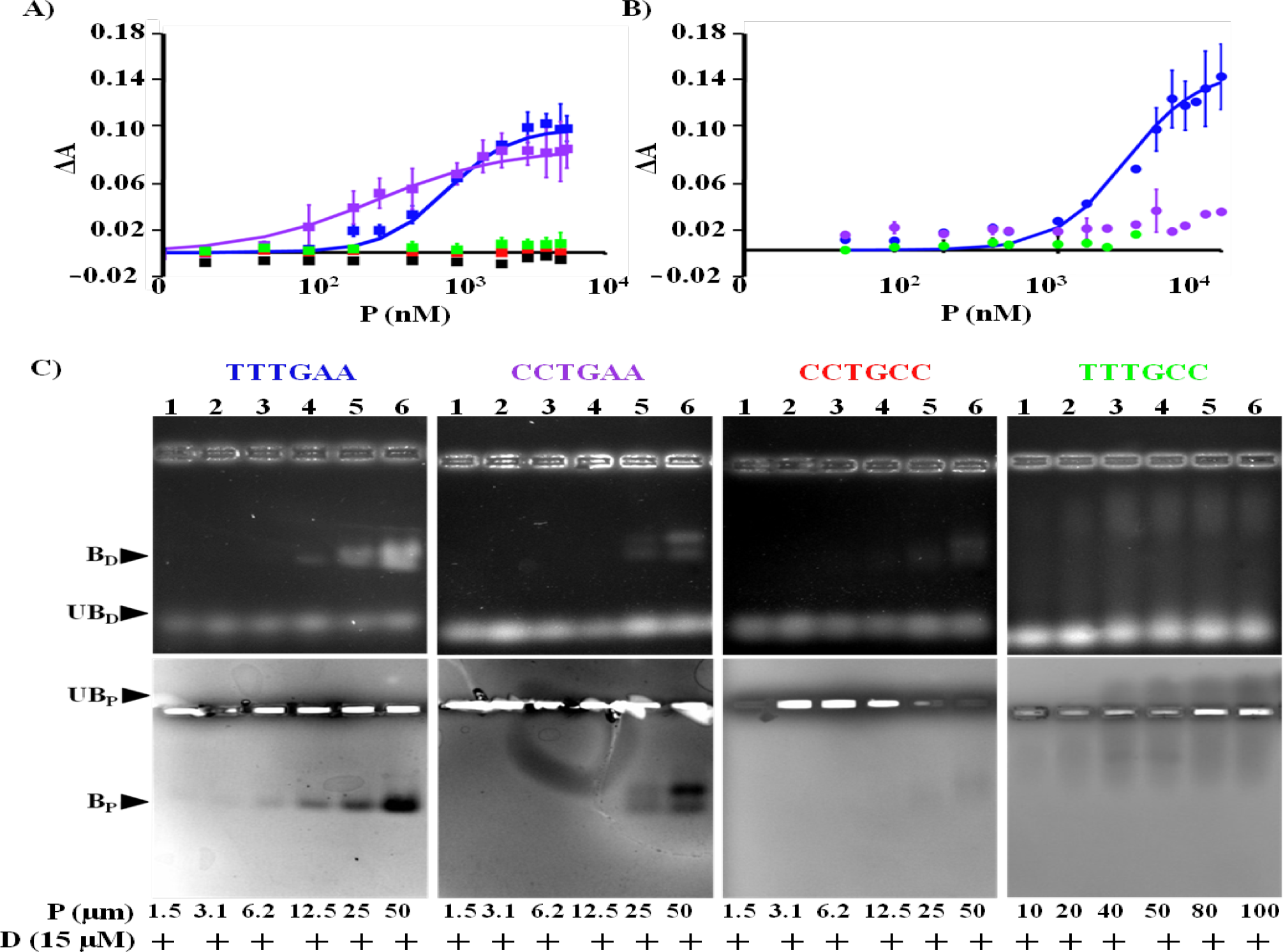
CHT7_CXC DNA binding specificity. **A, B)** Fluorescence polarization assay (FPA) demonstrating change in fluorescence anisotropy (ΔA) of 5nM 6-FAM labeled DNA with change in concentration of **A)** CHT7_CXC when DNA is 27mer duplex (5’-TATCTGGTGX^1^X^2^X^3^X^4^X^5^X^6^TTTCTGGATCTG-3’). Where X^1^-X^6^ is TTTGAA, CCTGAA, CCTGCC and TTTGCC are shown in blue, purple, red and green, respectively. The 13 mer duplex containing TTTGAA is shown in black. All data points are represented by squares and **B)** CXC_N when DNA is 27mer duplex containing TTTGAA, CCTGAA or TTTGCC, shown in circles in blue, purple and green, respectively. Where binding happens, the sigmoidal fit to ΔA is shown in bold line. Deviations shown are the standard errors estimated across the means of three different experiments. **C)** Electrophoretic mobility shift assay (EMSA) showing binding of CHT7_CXC to 27mer DNA duplex. Gel pictures in top row are the ones stained for DNA and in the bottom row corresponds to same gels stained for protein. Protein and DNA concentrations are mentioned at the bottom of the gels. Bound and unbound Protein and DNA are marked as (B_P_ and UB_P_) and (B_D_ and UB_D_), respectively.

### Differential binding abilities of CHT7_CXC’s subdomains impart sequence selectivity

The sequence-specific binding of CHT7_CXC towards TTTGAA seems similar to that of the LIN54. Hence, CHT7_CXC may also possess Tyr mediated binding, wherein, CXC_C should bind at T2 and T1*(A), and CXC_N should bind at A6 and A5*(T). In order to investigate the contributions of these subdomains in binding, the possible binding regions (T1T2 and A5A6) were individually substituted with the low-binding nucleotide pair (CC). This provides two variants of the binding region in 27mer DNA (5’-TATCTGGTG**CCTGAA**TTTCTGGATCTG-3’ and 5’-TATCTGGTG**TTTGCC**TTTCTGGATCTG-3’). Surprisingly, FPA shows efficient binding (K_D_ = 368.31±133.42nM, **Table1**) of CHT7_CXC towards CCTGAA (**Purple, Fig.1A**), while no binding for TTTGCC (**Green, Fig.1A**) containing sequences. These results on further validation through EMSA show that CXC binds CCTGAA in possibly two conformations as indicated by two bound fractions in Lane 5 and 6 of CCTGAA gels (**Fig.1C)**. Both the bound fractions of CCTGAA complex appear concurrently only after the protein was titrated beyond 12.5 μM. This, in comparison to the appearance of single bound fraction for TTTGAA at less than 6.2μM of the protein concentration (Lane 3, TTTGAA gels, **Fig.1C**), indicates that the two possible binding conformations of CXC with CCTGAA may possess approximately equal structural occupancies. CXC binds CCTGAA, likely through its CXC_N subdomain at its preferable binding site A6A5*, in spite of the absence of the binding site T2T1* for CXC_C. On the contrary, the appearance of smeared bound fractions of TTTGCC gels of **Fig.1C** indicates multiple weak binding conformations. Considering TTTGAA, both the subdomains have stable binding for their respective binding sites. The EMSA results for the variant sequences suggest that binding through CXC_N is necessary for binding of the CXC_C subdomain. The two bound conformations in CCTGAA complex probably results from the two different modes in which CXC_C may interact with the DNA, while the CXC_N remains anchored to the A6A5* nucleotide sites. This suggests that CXC_C induces an additional structural asymmetry in the CCTGAA bound conformations, in comparison to the TTTGAA bound conformation. Consequently, reduced depolarization of light is apparently reflected in the FPA using the CCTGAA sequence (**Fig.1A, Table 1**).

To understand, if the two subdomains possess independent binding abilities, the set of binding assays (FPA and EMSA) were conducted by titrating CXC_N and CXC_C concentrations in the presence of DNA substrates containing TTTGAA and its variants. CXC_C shows no binding with any of the DNA substrates (**Fig.S5**). The results from FPA show that the CXC_N could bind (K_D_ = 2633.33±906.20nM, **Table1**) TTTGAA (**Blue, Fig.1B**). Although the binding is much weaker than the full CXC domain. Similarly, EMSA results also suggest binding of CXC_N towards TTTGAA and CCTGAA (marked by a dotted region in **Fig.S6 &** bands in yellow in **Fig.S8** CXC-N gel) containing DNA. While none of the assays showed any signatures of binding towards TTTGCC by either of the subdomains.

Next, to examine if the mere presence of individual subdomains in each other’s vicinity suffice their binding abilities, we conducted EMSA in presence of both the subdomains (not linked by the loop). We observed the appearance of smear in the binding assays with the TTTGAA and CCTGAA containing DNA substrates (**Fig.S7 &** CXC_N+C gel in **Fig.S8**). Which could be due to the dissociation of the complex during electrophoresis. When compared to the binding studies of the full CXC (CXC gel **FigS8**), that shows stronger binding affinities, the results suggest that the loop that links the two subdomains is necessary for the efficient DNA binding.

All together, the results from these studies indicate that the two subdomains of the CHT7_CXC possess different DNA binding abilities. Wherein, the CXC_C binding onto the DNA seems to depend upon the binding status of the CXC_N and imparts sequence specific DNA binding to CHT7_CXC.

### Structural aspects explain sequence-specific DNA binding of the CHT7_CXC subdomains

To understand the molecular mechanism behind DNA recognition by CHT7_CXC, its homology model (c-score of -1.03, TM-score 0.7932, and RMSD of 0.37Å with LIN54 (**20**)) was obtained. The structural model docked to the 12mer DNA sequence containing TTTGAA as the binding site (5’-GAG**TTTGAA**ACT-3’) provided a DNA-bound complex of CHT7_CXC in which the sequence-dependent features remained preserved in terms of the dinucleotide geometry. Further, on equilibration through MD simulations, the bound complex retained all the specific interactions, similar to that in LIN54. Each CXC subdomain contained three Zn ions, covalently coordinated in the tetrahedral geometry with nine cysteine’s. In TTTGAA complex, the binding region is recognized by two subdomains, CXC_N and CXC_C connected by a loop. Tyrosine residues present in each of the subdomains show Hydrogen Bond (HB) interactions (dashed lines in purple, **Fig.2A**) with the step nucleotides in the DNA minor groove. Tyr17 in CXC_N binds at A6 and A5*(T), whereas Tyr99 in CXC_C bind at T2 and T1*(A). The maintenance of these HBs during the simulation time (Blue, **Fig.2B**) reflects the conformational stability of the DNA binding pockets surrounding the binding regions in TTTGAA. However, when the six nucleotide binding stretch was replaced with CCTGCC, none of the CXC subdomains possessed any Tyr mediated HB interactions in the DNA minor grooves (Red). While for CCTGAA, CXC_N stably maintains the Tyr17 mediated HB’s with A6 and A5* nucleotides, whereas no Tyr99 mediated HB interactions were observed due to the CXC_C subdomain (Purple). On the other hand, in TTTGCC complex, none of the subdomain Tyrosine show any HB interactions in the DNA minor grooves (Green).

**Fig2:**
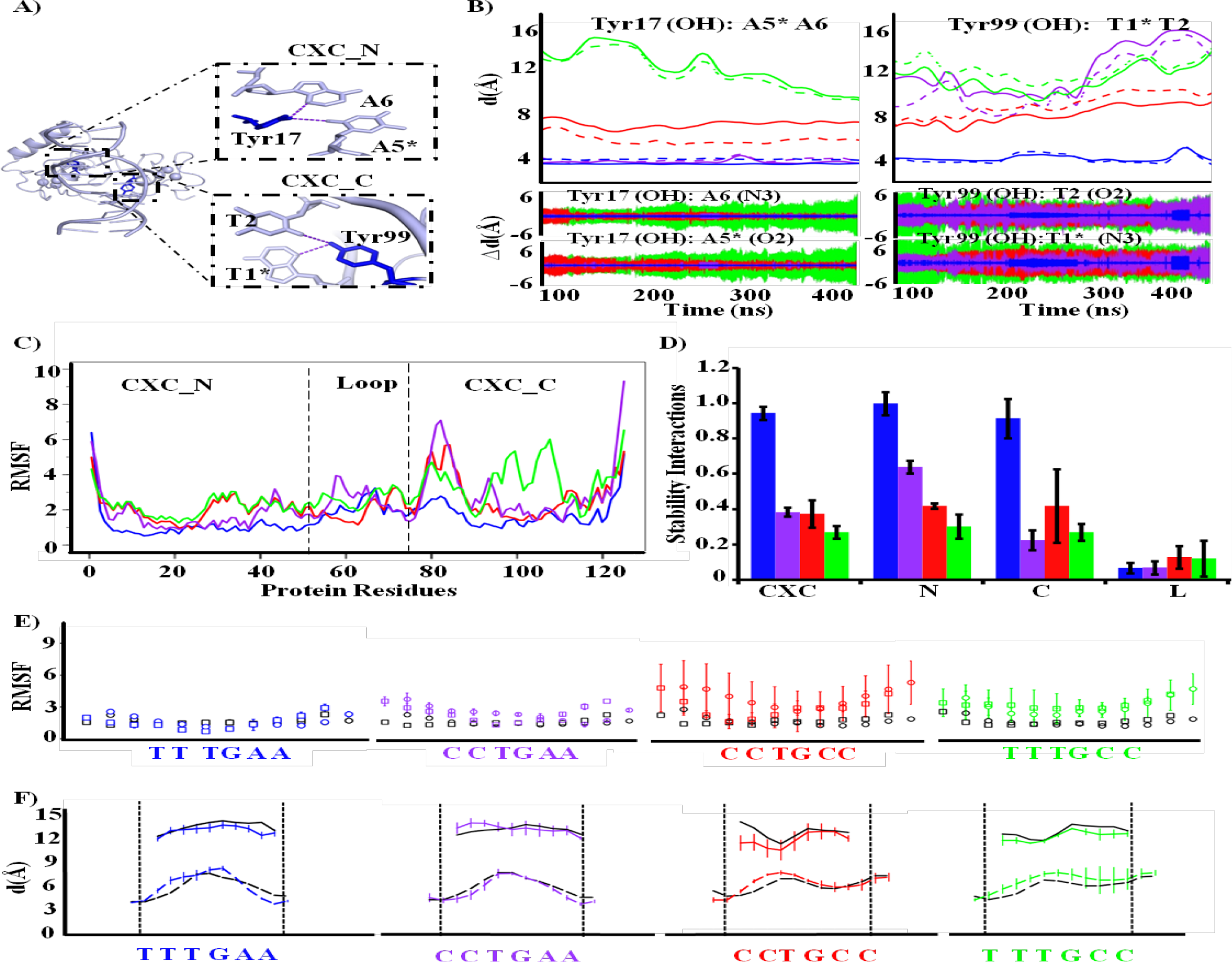
Structural aspects of CHT7_CXC and DNA complex. A) CHT7_CXC-DNA(with TTTGAA as binding site) complex structure obtained after 500ns MD simulation shown in cartoon. Tyr17 and Tyr99 (shown as sticks in blue) residues of CXC_N and CXC_C subdomains form H-bonds (purple dotted lines shown in zoomed in view) with A5*A6 and T1*T2, respectively. * represent complementary nucleotide. B) Averaged temporal variation of the distance (from three repeat simulations) corresponding to the Tyr mediated H bond distances (Left panel: Tyr17 and Right panel: Tyr99) in the complexes containing DNA. Protein C) RMSF and D) Stability interactions for different DNA sequences. E) RMSF of the phosphate atoms of DNA. 5’-3’ strand shown as open circles and 3’-5’strand shown as open square. F) DNA major (solid line) and minor (dotted line) groove widths. DNA in the bound complex (with TTTGAA, CCTGAA, CCTGCC and TTTGCC as binding regions, are color coded as blue, purple, red and green, respectively) and isolated DNA is in black. Error bars and deviation (Δd) are the standard errors from three repeat simulations

The tyrosine-mediated binding site recognition in the CXC subdomains, are interdependent upon two factors (i) the overall plasticity of the protein subdomains in the presence of the DNA sequence and (ii) the changes in DNA geometry in the bound complexes. Residue-wise Root Mean Square Fluctuations (RMSF) provide the signature of subdomain plasticity. CHT7_CXC and its CXC_N subdomain show relatively low RMSF in TTTGAA (Blue) and CCTGAA (Purple) bound complexes, in comparison to CCTGCC (Red) and TTTGCC (Green) complexes (**Fig.2C**). Further the subdomain plasticity was quantified as the stability interaction index which is the number of interactions (between the protein and the DNA within 4Å) formed per structure in a simulation (500 ns), given a prior possibility to exchange interactions from the pool of all the unique interactions for a complex. The stability interaction index was estimated for the CXC, its subdomains and loop for all the four protein-DNA complexes (TTTGAA; Blue, CCTGAA; Purple, CCTGCC; Red and TTTGCC; Green, **Fig.2D**). Higher stability index for the regions possessing low RMSF and vice versa shows concurrence between subdomain plasticity and their structural variations. Low structural variations between the first principal components (PC1) obtained from the simulated structures of CHT7_CXC DNA complexes suggest that CXC_N subdomain anchors onto the minor groove region surrounding the A5A6 position in TTTGAA and CCTGAA bound complexes (**Fig.S9**). Whereas, CXC_C stably surrounds the minor groove encompassing T1T2 in TTTGAA (Blue), while it loses its stability interaction in CCTGAA due to CC (Purple, **Fig.2D**, and **Fig.S9**). On the contrary, the unavailability of tyrosine-mediated binding sites for CXC_N at the 5^th^ and 6^th^ position in TTTGCC, destabilizes its overall binding (Green, **Fig. 2D**) and ultimately forces CXC_C, via the loop domain, to destabilize interactions with the DNA (**Fig.S9**). As a result, in TTTGCC containing DNA sequence, CXC_C residues high RMSF (**Fig.2C**) and reduced interaction stability (**Fig.2D**) hampers the subdomain’s ability to undergo tyrosine-mediated recognition in spite of the presence of its favorable binding region T1T2. Less stability interaction for the CHT7_CXC loop (**Fig.2D**) and comparatively longer (23 aa) loop size than that in LIN54 (14 aa) could impart extra flexibility to the subdomains in CHT7_CXC. PC1 from the LIN54 DNA complexes shows that structural variations due to the individual subdomains are less pronounced (black, **Fig.S9**). Swapping of the LIN54’s loop with that of the CHT7_CXC, in case of CCTGAA complex, enhances (Orange arrow, **Fig.S9**) the variations in PC1 (Grey, **Fig.S9**) due to CXC_C subdomain and indicates that the loop size could impact subdomain’s occupancy in the minor groove. Further, to test the impact of the loop on the distribution of the contacts (**SI**), we conducted Kolmogorov Smirnov (KS) test **(24)** on the set of different DNA complexes of CHT7_CXC, wild type LIN54, and modified LIN54 (LIN54 with its loop swapped with that of the CHT7_CXC loop domain). The results (**Table2**) indicate that except the CHT7_CXC’s complexes with TTTGAA and CCTGAA, contacts in all the other complexes possess significant (p value < 0.05) dependence on the loop.

**Table 2:** Kolmogorov Smirnov (KS) test showing p values to compare the significance of difference between the contact distributions of the sub domains in presence and absence of the loop.

| **Protein**<br>**→** | **CHT7_CXC** |  |  | **LIN54** |  |  | **LIN54 with CHT7's CXC**<br>**loop** |  |  |
| --- | --- | --- | --- | --- | --- | --- | --- | --- | --- |
| **DNA↓** | **CXC_N**<br>**+**<br>**CXC_C** | **CXC_N** | **CXC_C** | **CXC_N**<br>**+**<br>**CXC_C** | **CXC_N** | **CXC_C** | **CXC_N**<br>**+**<br>**CXC_C** | **CXC_N** | **CXC_C** |
| **TTTGAA** | **0.52**<br>**(3)** | **0.73** | **0.37** | **10<sup>-9</sup>**<br>**(1)** | **10<sup>-4</sup>** | **10<sup>-4</sup>** | **10<sup>-5</sup>**<br>**(2)** | **10<sup>-4</sup>** | **0.02** |
| **CCTGAA** | **0.97**<br>**(3)** | **0.11** | **0.12** | **<10<sup>-16</sup>**<br>**(2)** | **10<sup>-10</sup>** | **10<sup>-8</sup>** | **<10<sup>-16</sup>**<br>**(1)** | **10<sup>-11</sup>** | **10<sup>-7</sup>** |
| **CCTGCC** | **10<sup>-16</sup>**<br>**(3)** | **10<sup>-6</sup>** | **10<sup>-10</sup>** | **10<sup>-13</sup>**<br>**(2)** | **10<sup>-5</sup>** | **10<sup>-9</sup>** | **10<sup>-11</sup>**<br>**(2)** | **10<sup>-11</sup>** | **10<sup>-4</sup>** |
| **TTTGCC** | **10<sup>-8</sup>**<br>**(3)** | **10<sup>-4</sup>** | **10<sup>-4</sup>** | **10<sup>-14</sup>**<br>**(1)** | **10<sup>-9</sup>** | **10<sup>-6</sup>** | **NA** | **NA** | **NA** |

() parentheses represent number of simulation repeats

Next, the interdependence between the CXC_DNA complex formation and the DNA geometry was investigated. The preliminary observation comparing the RMSF of the phosphate atoms of the DNA in the bound complex with that of the DNA alone (Black, **Fig.2E**), illustrates that tyrosine-mediated binding has little effect on DNA in the TTTGAA complexes (Blue). Whereas, an increase in overall RMSF in CCTGCC (Red) and TTTGCC (Green) complexes signifies that weak or non-specific binding for these sequences increases the dynamics of the DNA backbone. While in CCTGAA complex, relatively higher DNA dynamics was observed at C1 and C2 positions than at A5 and A6. Exploring the DNA geometry, we observed a decrease in minor grooves width (dotted lines, Fig.2F) at the sites where the CXC subdomains bind (**Fig.2F**). This reduction, mainly at the pocket where CXC_N subdomain binds, is observed in the TTTGAA (Blue) and CCTGAA (Purple) bound complexes, in comparison to the unbound DNA (Black). The significant decrease (>3 standard deviations) at the A5A6 binding region, indicates that the CXC_N subdomain imparts higher binding stringency than the CXC_C subdomain. Whereas, increase in width or large deviations thereof represent either unbound subdomains, as in CCTGCC (Red) and TTTGCC (Green), or loose binding of CXC_C at T1T2 site of TTTGAA (Blue). Apart from this, changes were also observed in some of the base pair parameters of the DNA (**Fig.S10**). Relatively larger (∼15°) reduction in the axis bending was observed for the CHT7_CXC DNA complex having CCTGCC or TTTGCC than having TTTGAA or CCTGAA sites (∼4°). This difference is evident from the changes observed at the regions, T1T2 and A5A6. Similarly, a significant reduction (>4°) was also observed in the base pair inclinations of the DNA in the complexes having CCTGCC and TTTGCC. Whereas an average twist between the adjacent base pairs increases (>5°) in case of TTTGAA complex. Furthermore, the ionic dispersion around DNA provides additional validation regarding the asymmetric nature of CXC binding. In comparison to relatively uniform ionic concentration across the isolated DNA (dotted lines, **Fig.S11**), a sharp dip (Orange arrows) in concentration is found near the binding regions T1T2 and A5A6 in TTTGAA (Blue) and A5A6 in CCTGAA (Purple) complexes (Solid line). The reduction in ionic concentrations at the binding sites gets compensated by accumulated distribution (Grey arrows) around the T3G4 region in the DNA sequences.

Further on the conformational possibilities of all four CHT7_CXC-DNA (TTTGAA, CCTGAA, CCTGCC and TTTGCC) complexes were explored using Markov State Modeling (MSM) approach. **Fig.3** shows the possible number of metastable states and their occupancies. Predominantly single occupancy for TTTGAA (Blue) complex and two metastable states with equivalent occupancies for CCTGAA (Purple) complex supports our experimental observations (**Fig.1C**). The distribution of conformations for CCTGCC (Red) and TTTGCC (Green) is over three (34%, 26% and 37%) and four (17%, 14%, 28% and 41%) metastable states, respectively and reasons for why in the binding assays (**Fig.1A and 1C)** we observed weak and dispersive binding for these sequences.

**Fig3:**
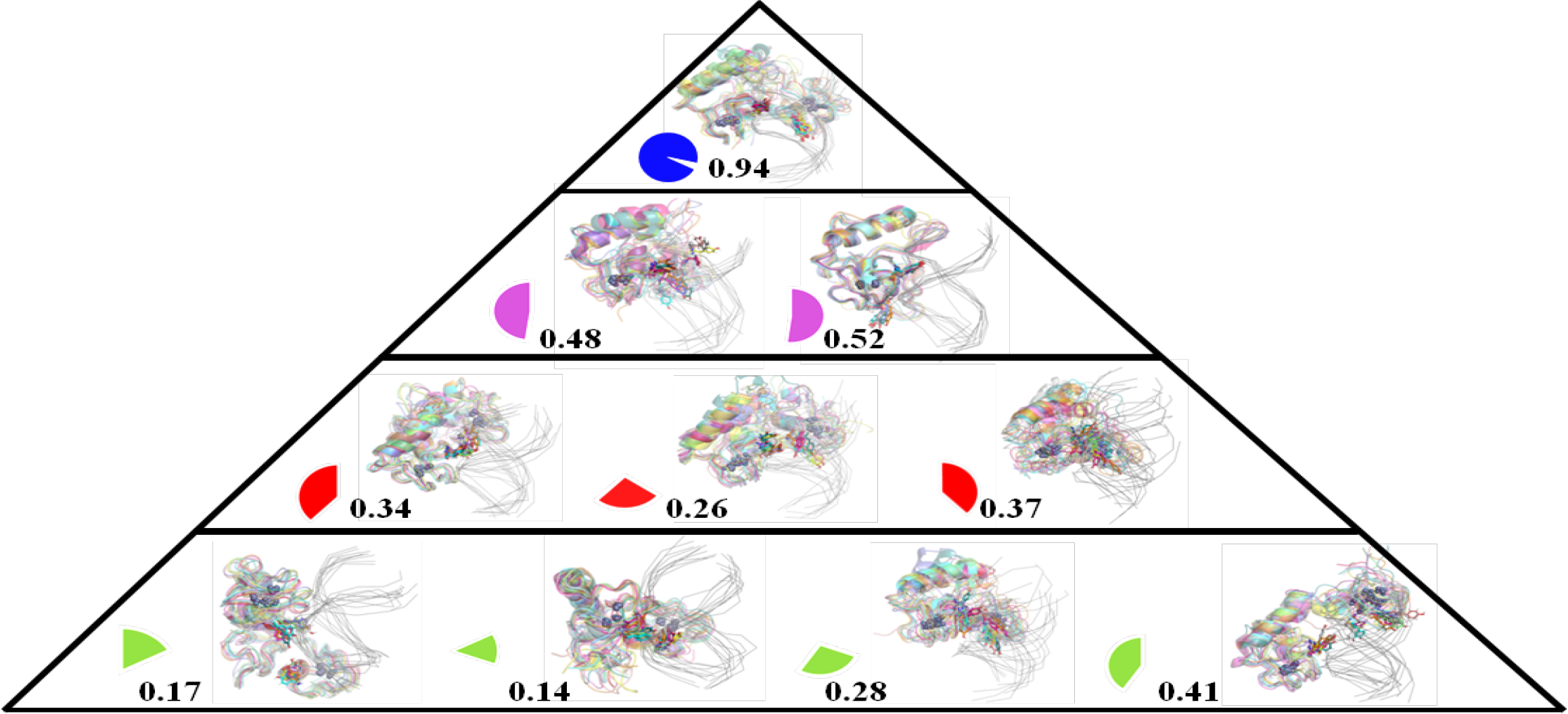
PCCA++ metastable states of CHT7_CXC DNA complexes. Shown are the metastable states that possess more than 10% occupancies. Complexes with DNA binding regions TTTGAA, CCTGAA, CCTGCC and TTTGCC possess 1, 2, 3 and 4 metastable states, respectively. The occupancies of the respective states are represented by the size of the pie (number depict fraction of the occupancy), color coded in blue, purple, red and green.

### Allosteric forces and not the space availability, stops CHT7_CXC from transcription activities

Using the CHT7’s alpha fold structural model **(25, 26)**, we estimated 100742.60 Å^3^ (**Fig.4A**, Brown sphere) of the enclosure volume within CHT7. This space is more than threefold of the net space (31179.98 Å^3^) actually required by the CXC-DNA (13mer TTTGAA sequence) complex. Furthermore, one of the most flexible models of the metastable states of the CCTGAA complexes, occupy to the maximum extent ∼60000.00 Å^3^ of the enclosure volume. CHT7 could therefore, adequately incorporate longer DNA sequences and also much flexible binding conformations. Thus, the possibility that the space availability within CHT7 could restrict CXC from DNA recognition is less favorable.

**Fig4:**
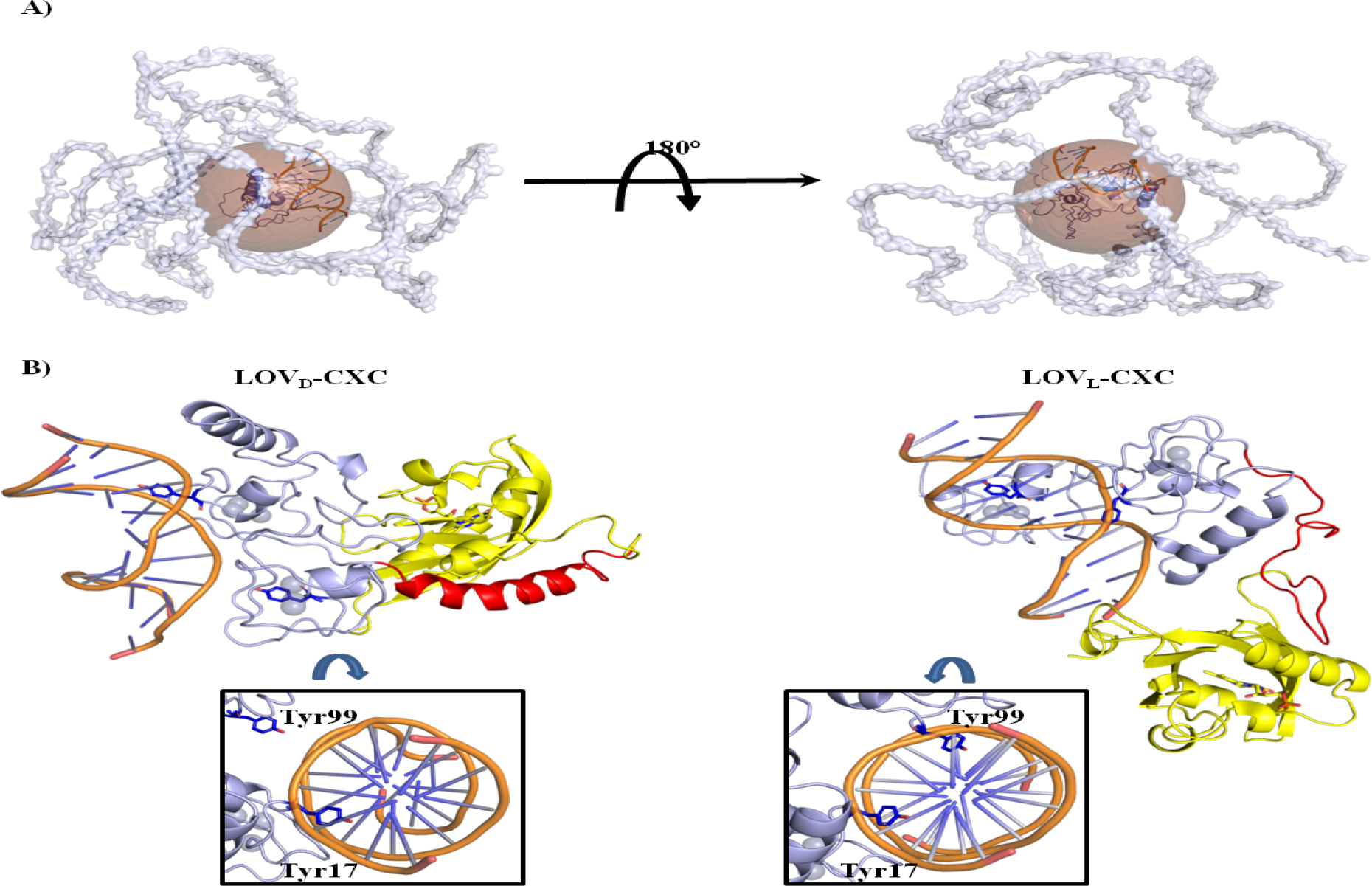
Spatial and allosteric restraints within CHT7. **A)** CHT7’s alpha fold structural model represented in the cartoon is shown from two different angles in grey. Enclosed within it is the brown sphere representing the inner boundary line of the CHT7 residues excluding and surrounding the CXC domain. Enclosed within this sphere is the CXC DNA (TTTGAA) complex structure in cartoon. **B)** Molecular dynamic structures of the molecular fusion of *Avena Sativa’s* LOV2 (AsLOV2) and CHT7_CXC generated under the dark (LOV_D_-CXC) and the light (LOV_L_-CXC) conditions in complex with 12mer DNA duplex containing TTTGAA as the binding region. Core LOV domain, its J_α_ helix and CHT7_CXC domain along with the coordinated Zn ions (shown in sphere) and DNA duplex (in brown) are shown as cartoon in yellow, red and cyan, respectively. Tyr residues that can mediate H bond interactions in the DNA minor groove pockets are represented as sticks in the element colour. Top view focusing DNA and the binding pocket regions show that Tyr99 of the CXC_C subdomain moves out of the DNA minor groove in the dark fusion LOV_D_-CXC, whereas remains within the pocket in light fusion LOV_L_-CXC

Further, we tested if structural changes induced by the formation of the CHT7 complex could affect the CXC’s binding to the DNA. For this, the allosteric scenario was simulated by designing two structural models LOV_D_-CXC and LOV_L_-CXC, generated through fusing and equilibrating the *Avena Sativa’s* LOV2 (AsLOV2) C-terminal to CXC’s N-terminal. Here, LOV_D_ and LOV_L_ represent AsLOV2 in its dark and light states, respectively. AsLOV2 under the dark conditions has its C terminal alpha helix (J_α_ helix) in the folded form, whereas on blue light exposure the J_α_ helix unfolds **(27, 28)**. This characteristic of AsLOV2 was exploited to understand if DNA binding ability of the CHT7_CXC can get impacted due to the allosteric effects. For this, both the dark and the light fusions were equilibrated with 12 mer TTTGAA DNA duplex. It was observed, in the LOV_L_-CXC fusion complex, the CXC_N and the CXC_C subdomains maintain the tyrosine-mediated DNA binding (**Fig.4B**, LOV_L_-CXC structure) and possess the binding enthalpy of -42.55 +/-5.62 kcal/mol and -41.68 +/-7.25 kcal/mol, respectively (**Table3**). Whereas, in LOV_D_-CXC fusion complex the Tyr99 of the CXC_C subdomain breaks H bonds with T1*T2 and moves out of the DNA minor groove (**Fig.4B**, LOV_D_-CXC). Also, the CXC_C subdomain possess significantly higher binding enthalpy in LOV_D_-CXC complex (∼0.28 +/-4.03 kcal/mole) than in the LOV_L_-CXC complex (-41.68+/-7.25 kcal/mol). And, LOV_D_-CXC (black) in comparison to LOV_L_-CXC (Blue) is found to be dynamically less stable **(Fig.S12A**) and shows increased RMSF (black) specifically in the CXC region (**Fig.S12B**). Additionally, a significant increase of ∼47 kcal/mol (ΔH_CXC+DNA_ = -98.20 +/-10.12 kcal/mol to ΔH_LOVD-CXC+DNA_ = -52.06 +/-7.45 kcal/mol) in the binding enthalpy of the CXC has been observed in the dark fusion. These observations suggest that the folded state of the J_α_ helix in LOV_D_-CXC, in contrast to its unfolded state in LOV_L_-CXC complex, imposes steric restrictions on the CXC and refrains it from maintaining the stable binding with the DNA.

**Table 3:** Binding energies for Protein DNA (with TTTGAA binding site) interaction using GBSA.

|  |  |  |  |
| --- | --- | --- | --- |
| **(LOV+CXC)-DNA** | **CXC** | **NA** | **NA** |
|  | **LOV<sub>L</sub>-CXC** | **-159.62** | **-106±11.06** |
|  | **LOV<sub>D</sub>-CXC** | **-161.79** | **-51.75±7.47** |
| **(CXC)-DNA** | **CXC** | **-140.55** | **-98.20±10.12** |
|  | **LOV<sub>L</sub>-CXC** | **-156.03** | **-83.04±8.77** |
|  | **LOV<sub>D</sub>-CXC** | **156.49** | **-52.06±7.45** |
| **(LOV+CXC_N)-DNA** | **CXC** | **NA** | **NA** |
|  | **LOV<sub>L</sub>-CXC** | **-157.20** | **-66.38±8.71** |
|  | **LOV<sub>D</sub>-CXC** | **-160.26** | **-51.51±6.44** |
| **(CXC_N)-DNA** | **CXC** | **-134.17** | **-50.57±7.11** |
|  | **LOV<sub>L</sub>-CXC** | **-147.05** | **-42.55±5.62** |
|  | **LOV<sub>D</sub>-CXC** | **-149.29** | **-51.85±6.40** |
| **(CXC_C)-DNA** | **CXC** | **-130.94** | **-48.38±5.62** |
|  | **LOV<sub>L</sub>-CXC** | **-144.35** | **-41.68±7.25** |
|  | **LOV<sub>D</sub>-CXC** | **-140.32** | **0.28±4.03** |

All units are reported in kcal/mol.
GBSA calculations were carried out at 298K

## Discussion

In *C*.*reinhardtii*, CHT7 is a transcriptional regulator essential for the entry and the exit from the quiescence and successful progression of these cellular states, and for the accumulation of the TAG under the nutrient deprived conditions (3). CHT7 functions by forming large protein complexes, using its predicted protein binding domains (PPDs) to interact with the external transcription factors (7). Whereas, its DNA binding CXC domain remains effectively dispensable for the CHT7’s transcriptional activities. However, it is seen that several CXC domain containing proteins undergo their transcriptional activities via these domains (11–19). The present study demonstrates that the CXC domain within CHT7 is an asymmetric domain with abilities to efficiently scan DNA sequences, and could therefore interfere with the precise regulation of quiescence. Further, this study provides a mechanistic insight on how algal cells may regulate the DNA binding ability of the CHT7_CXC domain and switch them into an inactive state during the functioning of the CHT7 complexes.

CHT7_CXC’s close homolog LIN54 shows sequence-specific recognition for the TTTGAA stretch in the DNA **(20)**. The availability of this stretch in the promoter region of some of the genes that get misregulated in the cht7 mutant (**Fig.S3**) provided a basis to examine the CHT7_CXC’s DNA recognition abilities. We observed that CHT7_CXC shows sequence-specific binding affinity (K_D_ = 873.38±133.17nM) for 27mer DNA fragments containing TTTGAA region (**Fig.1A and 1C**). Further, our MD simulation studies suggested, like LIN54, CHT7_CXC also binds onto the TTTGAA stretch via its CXC_C and CXC_N subdomains (**Fig.2A**). These subdomains interact with the minor grooves of the DNA by forming Tyr mediated H Bond interactions with the T1*T2 and A5*A6 nucleotide pairs of the binding stretch, respectively. However, experimentally we couldn’t observe any binding when the shorter DNA fragment (13mer) was used (Black squares, **Fig.1A**), indicating that CXC binding is dependent on the DNA length. Possibly, the CXC undergoes a slide and search mechanism to recognize the specific binding stretch of the DNA. The longer DNA fragments (27mer) provide a platform containing the sites to which the CXC subdomains can transiently interact, to initiate the search for the specific target sites. The unavailability of sufficient sites, in a shorter DNA fragment, reduces the propensity of protein DNA encounter. Thus, the overall binding affinity of the CXC reduces for the shorter DNA fragments even though the specific binding stretch is available.

CHT7_CXC loses its binding ability if CXC_N specific binding region (A5A6) or if both the subdomain binding sites (A5A6 and T1T2) in TTTGAA are replaced with CC (Red and Green, **Fig.1 & Fig.1CA**). Whereas, a similar replacement only at the CXC_C specific binding site (T1T2), does not impair CXC from DNA binding (magenta, **Fig.1A & Fig.1C**). This indicates that the two subdomains possess asymmetric DNA binding abilities. This differential binding of the two subdomains is also reflected from the attributes like, differences in the plasticity possessed by these subdomains’ for different DNA stretches (**Fig.2A-D**), and variations observed in the geometric parameters (**Fig.2E, Fig.2F and Fig.S10**) and the ionic strength (**Fig.S11**) in the DNA. It was also observed that, in comparison to no binding by an isolated CXC_C (**Fig.S5 and Fig.S8**), CXC_N shows binding, although weak (K_D_ = 2633.33±906.2 nM), for the A5A6 region (**blue, Fig.1B, Fig.S6 and Fig.S8**). Whereas, the full CXC binds efficiently (K_D_ = 368.31±133.4 nM, **magenta, Fig.1A**) to the DNA substrate containing the CCTGAA. This shows that the binding efficiency enhances when the CXC_N is connected via a loop to the CXC_C. While the absence of the CXC_C’s specific binding sites (T1*T2), in CCTGAA, merely distributes the CXC_C occupancy on the DNA (**CCTGAA (magenta) in Fig.1C and Fig.3**). Whereas, absolutely no binding was observed for the DNA substrate containing TTTGCC, indicates, for the CXC_C to bind onto the T1*T2, CXC_N must be anchored at the A5*A6 site. We therefore expect that the CXC_C subdomain, due its low binding affinity within CXC, may essentially be utilized in scanning the potential binding sites from the pool of the non-specific sites in the *C*.*reinhardtii* genome.

Further, our results also suggest that the DNA recognition ability of CXC is partly dependent on the loop between the two subdomains. The overall binding stability, as is projected by the distribution of the contacts between the protein and the DNA (**Table2**) could depend upon two factors (i) the contact distribution of the individual subdomains and (ii) the spatial distribution of the loop (associated with the size of the loop). It was observed that in spite of the comparatively longer loop (in comparison to that in the LIN54), the overall binding of the CHT7_CXC onto the TTTGAA or the CCTGAA stretches, is little dependent on the loop (**Table 2**). This is because, the comparatively narrower distribution of the contacts due to the bound subdomains in these complexes, dominate over the unstability imparted due to the longer loop. Whereas, the loop’s flexibility and the broader distribution of contacts due to the weakly bound subdomains, synergistically induces reduced binding stability in case of the nonspecific and semi specific regions like CCTGCC (Red) and TTTGCC (Green), respectively (**Fig.S9**). While, on the other hand, the limited flexibility imposed by the shorter loop in LIN54 restricts the movement of the individual subdomains from possessing independent DNA recognition abilities. Possibly due to this reason, the LIN54 shows DNA binding only when the consensus sequence (TTYRAA) has both the T1T2 and the A5A6 in place (20). Whereas, in CHT7_CXC the subdomains are relatively flexible to recognize binding stretches other than the TTYRAA. Thus, the CHT7_CXC could bind and differentiate between the semi-specific stretches. Such as, it can bind at CCTGAA and not at TTTGCC.

This differential binding ability of CHT7_CXC’s subdomains is especially important in the context of *C*.*reinhardtti*, due to its GC-rich genome. The CHT7_CXC subdomain-specific binding sites (T1T2 and A5A6) are less abundant (Blue, **Fig.S3**) than the non-specific sites (CC) (Red, **Fig.S3**). Symmetric subdomains are found efficient for DNA binding, but show compromised DNA recognition kinetics (29). Whereas, asymmetry allows to conduct quick search for the specific binding sites from the pool of non-specific or semi-specific sites (29, 30). It also aids in the dissociation from the non-specific sites. Asymmetry between the tethered subdomains facilitates DNA search through the monkey bar mechanism in which different subdomains can scan distant DNA segments (29, 31). Egr-1 (30), Pax6 (32), Oct1 (32), Pax5 (33), and PARP1 (34) are some of the several DNA binding proteins that exploit their subdomain asymmetry to rapidly search specific binding regions in an ensemble of densely populated non-specific binding regions. We propose that CHT7_CXC could similarly utilize its subdomain asymmetry to quickly recognize low abundant specific (TTTGAA) and semispecific (CCTGAA) DNA binding sites. A previous study by Takeuchi *et al*, showed that together with the proteins involved in DNA metabolism, chloroplast division and the cell cycle regulatory kinases, other CXC domain containing proteins of *C*.*reinhardtii* also remain repressed in the presence of CHT7 and only get derepressed in the *cht7* mutant **(7)**. All together this indicates, to allow precise transcriptional regulation via the CHT7 complex, the *C*.*reinhardtii* cells keep all the possible cell cycle regulators under the check. It is therefore essential that the CHT7_CXC also remains effectively dispensable for any regulatory activities.

One possible way by which the CHT7 can restrict its CXC domain from DNA recognition is by imposing spatial constraints on this domain from undergoing DNA binding. We therefore explored if CHT7 has enough space available to accommodate the most flexible DNA bound CXC complex. Interestingly, it seems that a sufficiently big pocket (100742.60 Å^3^) is available within CHT7 to accommodate the CXC-DNA binding volume (**Fig.4A**). Therefore, the possibility that space within the CHT7 limits its CXC domain from DNA binding, can be excluded. It has been observed that several transcription factors (TF) that possess DNA binding for the specific set of consensus sequences in-vitro, do not exhibit a similar binding profile in-vivo (35). In vivo, the DNA binding specificity of the archetypal zinc finger TF’s (KLF3) DNA binding domain (DBD) strongly relies on its non-DBD. The non-DBDs regulate the binding specificity of the DBDs by either directly recruiting external TF’s or by binding to the cofactors that can direct external TF’s to the promoter sites and could therefore attain a different DNA binding profile than in vitro. In vivo, the TF could show specificity towards fewer (or no) binding regions than in vitro or may even show binding for absolutely new DNA sequences. We could therefore equate the DNA binding profile of any TF to be in parts due to, its in vitro binding profile, binding contribution due to the non DBD related factors (such as the external TFs or the cofactors), and the mutual effect of the individual DBDs of the parent TF and the external TF’s. This mutual effect could be a result of the conformational changes that DBDs may undergo on macromolecular crowding in a large protein complex (36) that is an assembly of the parent TF, external TFs, and the cofactors. The parent TF could therefore exploit this effect to regulate the binding efficiency of its DBD. We hypothesize that molecular crowding within the protein complex could bring conformational changes in CHT7, which may allosterically impact the DNA binding capabilities of the CXC domain. To capture the essence of the allosteric effect on CXC, we simulated the simplistic models in which the AsLOV2 was fused to the CHT7_CXC’s DNA bound state. Two different conformational states of the AsLOV2 were used (i) the dark state, here the C terminal J_α_ helix remains folded and could therefore impose steric forces on the CXC domain, and (ii) the light state, in which the J_α_ helix remains unfolded. The dark state situation mimics the molecular crowding due to the formation of the CHT7 complex and the light state corresponds to a situation in which CHT7 remains isolated. It was found that in the dark state fusion, the Tyr 99 of the CXC_C subdomain moves out of the DNA minor groove (**Fig.4B**). Also, the energetics (**Table3**) indicate that AsLOV2 in the dark state imposes steric forces on CXC, making it to unbind DNA. Whereas, the light state fusion favors the DNA binding of the CXC domain. It is therefore suggested that the molecular crowding due to the formation of the CHT7 complex could induce structural changes within the CHT7, that switches CXC to the conformational state incapable of DNA binding. Knowing that the CHT7_CXC domain is an asymmetric domain, such an allosteric switching mechanism could rapidly restrict this domain from interfering with the CHT7’s quiescence repression and accumulation of TAG. This mechanism could also be a much general approach, using which the TFs regulate the activities of their DBDs.

## Material methods

Promoter Prediction and CXC binding sites: Details are provided in SI.

Cloning and purification: Details for obtaining CHT7_CXC, CXC_N and CXC_C proteins provided in SI.

Binding Assays: Protein DNA binding was performed using FPA and EMSA. Details for which are provided in SI.

Computational approaches: Details of all the computational approaches viz Homology modelling, Protein and DNA docking, MD simulations, MSM studies and preparation of fusion models are provided in SI.

## Acknowledgments

We thank Professor. Deepak T Nair from Regional Centre for Biotechnology to allow MC to work in his laboratory to conduct a part of cloning, protein purification and DNA binding experiments. We are also thankful to Professor. Neel Sarovar Bhavesh from ICGEB, New Delhi for allowing MC to use their research facility for carrying out some protein purifications. AS acknowledge the funding support from his UGC-FRP Startup grant and the Research grant (BT/PR34567/BRB/10/1822/2019) from Department of Biotechnology.

