## Supplementary material for "A possible mechanistic insight on how Compromised Hydrolysis of Triacylglycerol 7 (CHT7) restrains the involvement of it’s DNA binding CXC domain from quiescence repression": supli.pdf

**Identification of promoter regions and CXC binding sites:**
53 photosynthetic, 250 flagellum, 8 CDKs and 2 autophagy genes have been identified as the misregulated genes in the *cht7* mutant (1). The phytozome13 database and NCBI's Genome viewer were used to acquire sequences for these genes. Two approaches were used to identify promoter regions. i) Promoter prediction using the PromPredict tool (2, 3) and (ii) 1000 bps upstream of 5'UTR was selected as the region containing the promoters.

**Cloning**: The codon-optimized CHT7's CXC domain gene (*CHT7_CXC*) was synthesized for the expression in *E.coli* (GenScript Biotech). The *gene* was inserted into the pGEX-6P1 (GE Healthcare) expression vector between the *BamH1* and *Xho1* restriction sites. Lane3, **Fig.S13** shows the double-digested plasmid. Since the inserted gene (*CHT7-CXC* gene) is only 384 bp, we couldn't see the insert band on the gel. The presence of the insert was therefore confirmed through sequencing. The two smaller gene fragments of the CXC N terminal (*CXC_N*) and the C terminal (*CXC_C*) were amplified from the *CHT7_CXC* gene and were cloned into the pGEX-6P1 vector.

**Protein purification**: The protein purification methodology adopted for CHT7_CXC was similar to as described in our earlier works (4, 5). The expression of CHT7_CXC (123 bp) along with the GST tag was standardized in *E.coli*, Rosetta DE3 cells. For purification, 6 liters of LB media was inoculated with 50 ml of primary culture at 37°C. When OD reached 0.6, the culture was induced with 0.4mM IPTG, and 500 µM $ZnSO_4$ was added. The culture was incubated for 18 hours at 18°C. Cells were pelleted down at 6000 rpm for 15 minutes and resuspended in lysis buffer ( 25 mM HEPES buffer pH 7.5, 5 mM β- mercaptoethanol, 1mM PMSF, 250mM NaCl, 0.1% Triton X100 and 10% Glycerol). Resuspended cells were sonicated and centrifuged at 15000 rpm for 1 hour. The filtered supernatant was passed through the GST-Sepharose column (GE Healthcare). GST tag was then removed using PreScission protease (GE Healthcare). The purity of the CHT7_CXC protein was assessed by SDS–PAGE (Lane 3, **Fig.S14**). The concentration of protein was estimated by BCA assay using Thermo Scientific kit. Mass spectrometry was used to confirm the molecular weight (13.86 kDa) of CHT7_CXC. CHT7_CXC was further purified through gel-filtration chromatography using the Superdex 75 column (GE Healthcare). No significant peak, from the gel filtration, could be observed due to the lack of Tryptophan residues within CHT7_CXC. The eluate was therefore collected in fractions of equal volumes and run over the SDS page to confirm the presence of the protein. Approx ~3 to 5 mg of protein from 6 liters of the starting cultures could be obtained. The sequence of the purified CHT7_CXC was also confirmed using Mass spectrometry. CXC_N (7 kDa) and CXC_C (4 kDa) subdomains were purified using a similar protocol. The subdomain protein used for the binding assays was purified until the GST column purification step.

**Fluorescence Polarization Assay (FPA):**

The 6-carboxyfluorescein (6-FAM) labeled DNA was titrated with varying concentrations of purified protein. DNA (5nM) was titrated against different protein concentrations from 0 to 10 μM. Buffer composition in all the experiments was HEPES 25mM, beta-mercaptoethanol 5mM, PMSF 1mM, NaCl 100mM, Glycerol 2%. Assays were conducted using the multimode plate reader, M5 Spectramax (Molecular Devices) as done previously (6)**.** The fluorescence anisotropy values were acquired by exciting the sample at 495 nm and emission was recorded at 517nm both in parallel ($I_{par}$) and perpendicular ($I_{per}$) polarization. Three independent readings were taken for each protein concentration. Reduced anisotropy values were calculated by subtracting the value of the reaction without protein from the anisotropy values of the reactions with protein. The degree of depolarization of light (i.e. anisotropy) was calculated as follows:

$A = (I_{par} - G*I_{per})/( I_{par} +2G*I_{per})$, where G (G factor)= 1.000

Reduced anisotropy with the protein concentration was modeled using the Hill1 equation:

$Y = A_{min} + (A_{max} - A_{min}) [X^n/(K_d^n+X^n)]$,

where X is the concentration of protein, Y is the reduced anisotropy, $A_{min}$ is minimum reduced anisotropy, $A_{max}$ is maximum reduced anisotropy, $K_d$ is protein-DNA dissociation constant and n is the Hill coefficient.

**Electrophoretic Mobility Shift Assay (EMSA)**:

The reaction mixtures were prepared by titrating protein concentration up to 100 µM against 15µM of ds DNA oligomers. Other reaction components used were 25 mM HEPES pH 7.5, 5 mM β- mercaptoethanol, 1mM PMSF, 250mM NaCl, 0.1% Triton X100 and 10% Glycerol. Reaction mixture was incubated for 1hr at $4^oC$ and was run in 0.6% Agarose gel with Tris-Boric acid at pH 6.8 at 35 Volt. DNA was visualized through EtBr staining and protein through Brilliant Blue (Sigma-Aldrich) stain (7).

The sequences of the DNA oligomers used were:

27mer TTTGAA: 5’-TATCTGGTG**TTTGAA**TTTCTGGATCTG-3’
27mer CCTGAA: 5’-TATCTGGTG**CCTGAA**TTTCTGGATCTG-3’
27mer CCTGCC: 5’-TATCTGGTG**CCTGCC**TTTCTGGATCTG-3’
27mer CCTGCC: 5’-TATCTGGTG**TTTGCC**TTTCTGGATCTG-3’
13mer TTTGAA:5’-GGTG**TTTGAA**TTT-3’
29mer without any TTYRAA site: 5’ GATCAATAGATGCGCAGATCATTTACATT 3’
LHCA6 DNA substrate: 5’-TGGACTGGATTTGAACGATTGAAAGCT-3’
LHCB5 DNA substrate: 5’-TTCTTTTCATTTGAATGGTTTTCCGAC-3’

To be noted, the protein used for LHCA6 and LHCB5 EMSA runs was GST column purified. Therefore, the bound fractions appeared as a smear (Fig.S4), unlike the single band in case of 27mer TTTGAA (Fig1.C) where the protein used was further purified by gel filtration chromatography after the GST column purification.

#### Homology Modeling of protein and DNA

A set of five homology models for CHT7_CXC using LIN54 (PDB ID: 5FD3, (8)) as a template was obtained using the advanced modeling techniques of I-TASSER suite (9). Of these models, one with the best confidence score (c-score), TM-score, and lowest RMSD with reference to LIN54 structure was selected for further studies. Similarly, a homology model was also obtained for LIN54 with its loop swapped with that of the CHT7_CXC. Whereas, homology models for 12 mer DNA sequences with four stretches TTTGAA, CCTGAA, TTTGCC and CCTGCC, were modeled using 3D-NuS (10) by taking DNA in 5FD3 (8) as a template.

#### Protein and DNA docking

Protein DNA docking was conducted using HADDOCK 2.4 (11, 12) . The distance restraints were applied to retain key interactions known from the human LIN54-DNA complex (PDB ID: 5FD3) and to retain classical tetrahedral coordination for Zn and Cysteine’s. The major challenge was to dock the CXC domain with DNA sequences containing CCTGCC, CCTGAA and TTTGCC as binding sites as their interactions were not known. However, we aimed to retain all the essential sequence-specific interactions and other nonspecific interactions based on the known physico-chemical properties like shape complementary, electrostatics and scoring obtained from the docking software. Finally, before taking the best model as the starting structure for molecular dynamic simulations, torsion adjustments were made using Chimera (13).

#### Molecular Dynamics simulations of CXC-DNA complex

The MD simulations and binding energy calculations in this study were carried out using MD simulation program AMBER (14). Set of three MD simulations (500 ns production run) were carried out on all the docked protein DNA complexes as the starting structures. The complexes of the proteins viz CXC, CXC_N, CXC_C, LIN54 and LIN54 (with swapped loop) with the

12mer DNA (GAGX1X2X3X4X5X6ACT) sequences having X1-X6 as TTTGAA, CCTGCC, CCTGAA and TTTGCC, were studied.

All MD simulations were performed using the AMBER ff14SB and bsc1 force fields. Complexes were solvated in a 10 Å isometric box containing TIP3P water molecules. $Na^+$ ions were then added to neutralize the system and the solvent concentration was made to 150 mM of NaCl by adding an adequate number of $Na^+$ and $Cl^-$ ions in the box. The resulting solvated system was energy-minimized in three steps. First, both protein and DNA were fixed using a positional restraint on each of the atoms to its starting structure with a force constant of 500 kcal $mol^{-1}$ $Å^{-2}$, and the position of the water and ions were allowed to change using 1500 steps of steepest descent minimization, followed by 1500 steps of conjugate gradient minimization. The second minimization was done using a force constant of 25 kcal $mol^{-1}$ $Å^{-2}$. The whole system was then energy-minimized for 5000 steps, including 2500 steps of steepest descent minimization. In the final minimization step, the entire system, containing a DNA protein complex with ions and water molecules was subjected to 5000 steps of minimization without any restraints and another 2500 steps of steepest descent minimization. The system's temperature was then raised from 0 to 300 K at constant volume for 20 ps, while the protein and DNA complex were weakly restrained at 10 kcal $mol^{-1}$ $A^{-2}$. For the next 20 ps the system was subjected to the Langevin temperature equilibration scheme (NTT =3) and SHAKE constraints were applied to all bonds involving hydrogen. Then the restraints on protein and DNA were relaxed to 0.5 kcal $mol^{-1}$ $A^{-2}$ and the system was equilibrated at constant volume for 80 ps. Finally, the 500 ns production simulations were carried out at constant temperature (300 K) and pressure (1 bar) with a 2 fs time step.

**Impact of the loop on the distribution of the protein DNA contacts:**

The impact of the loop was estimated on the distribution of contacts formed between the protein subdomains and the DNA. The interatomic distance less than 4Å was considered as a contact. The distribution of the contacts was calculated using all the structures derived from the MD simulations of the protein DNA complex. Two kinds of complexes were taken into account, (i) that included and (ii) that excluded the loop between the protein subdomains. The KS (Kolmogrov Smirnov) test was then applied to measure the significance (p value, **Table 2**) of the difference between the distributions estimated using both the kinds of the complexes. For any protein DNA complex we estimated this significance value for three cases, where the contacts were considered due to (i) both the subdomains (CXC_N+CXC_C), (ii) only CXC_N and (iii) only CXC_C. This exercise was carried out using the structures from the MD simulations of all the 12 combinations of the protein DNA complexes formed using the three different proteins (CHT7_CXC, the LIN54, and the LIN54 in which its loop was swapped with the CHT7_CXC's loop) and four DNA sequences having TTTGAA, CCTGAA, TTTGCC and CCTGCC as the binding regions. Number of replicates of each simulation is mentioned in the parenthesis in **Table 2**.

**Markov State Modelling (MSM)**

The structural data was gathered from atleast three independent MD simulation repeats of 500ns each for any protein DNA complex to perform MSM (15) analysis using Pyemma (version 2.5.7, (16)). Equilibration dynamics of these simulations suggested that the data used is sufficient to capture the slowest dynamical features and to scan the complete conformational space. To reduce the high-dimensional phase space, H-bond distance between the protein and the DNA was found to be the most relevant feature to effectively capture the slowest components in the structural changes. Only those H-bonds were considered that appeared in atleast 75%, 60% and 50% of times in each of the simulation replicates of the TTTGAA, CCTGAA and CCTGCC complexes, respectively. The TTTGCC trajectories did not show a sufficient % of common H bonds between the replicates, three of the most frequently appearing H-bond distances in each replicate were considered as the feature for the dimensionality reduction. The complete conformational phase space could be captured through the Time Independent Component Analysis (TICA) (17, 18) using only 19, 29, 19, and 37 degrees of freedom for TTTGAA, CCTGAA, CCTGCC, and TTTGCC complexes, respectively. The free energy distribution maps (**Fig.S15A**) were produced by the projection of the transitions in the two slowest TICA components (IC1, IC2). The TICA coordinates were clustered into various discrete states using the k-means approach (19). State discretization was carried out using different numbers of cluster centres (5, 10, 15, 20, 30, 75, 200, 450). The VAMP-2 score got saturated after 200 cluster centres for TTTGAA and CCTGAA whereas for CCTGCC and TTTGCC after 100 and 30, respectively. The estimated implied timescales (its, **Fig.S15B**) and Chapman-Kolmogorov test (20) were then used to validate the lag time. Bayesian MSM models were generated using the lag time of 6, 10, 3 and 8 steps for TTTGAA, CCTGAA, CCTGCC and TTTGCC respectively, wherein the step size is 0.02ns. Number of the metastable states in the MSM study for each complex was selected by considering the highest time scale separation between the eigenstates (**Fig.S15C**). Using the PCCA++ (21), we could resolve these metastable states within the first two TICA components. Each metastable state is represented by 10 models (**Fig 3**).

**Volume estimation**

The enclosure volume within CHT7 that can accommodate CXC-DNA complexes (Brown sphere, Fig 4A) was estimated using vmd extension epock (22). Enclosure is marked by the inner lining of all the residues that surround the cavity in the CHT7 alpha fold structure, after excluding the CXC domain from the protein. And, the net pocket volume of the CXC_DNA complex was estimated using $V_{net} = V_{CXC} + V_{DNA} – V_{DBP}$. Here, the $V_{CXC}$, $V_{DNA}$ and $V_{DBP}$ are the volumes of the sphere that can enclose the CXC domain, the cylinder that can enclose the DNA, and the sphere that can enclose the DNA binding pocket, respectively. These were estimated to be 26279.00 $Å^3$, 9065.00 $Å^3$ and 4164.12 $Å^3$, respectively for the 12 mer CXC_DNA (12 mer, 5'-GAGTTTGAAACT-3') complex.

**Preparation of LOV_CXC fusion models**

$LOV_D$-CXC and $LOV_L$-CXC are the respective dark state and the light state fusions of the AsLOV2 and the CHT7_CXC. $LOV_D$ is a crystal structure of AsLOV2 (PDB ID: 2V1A, (23)) in the dark state, in which the C-terminal $J_\alpha$ helix is in folded conformation. Whereas, $LOV_L$ is the AsLOV2's light state structure showing a completely unfolded $J_\alpha$ helix. $LOV_L$ was obtained through simulating the light induced condition modeled into the dark structure 2V1A. The complete methodology of this simulation is explained below in the section "Generation of $LOV_L$ conformational state". Fusions were modeled using an extensible molecular modeling system CHIMERA (13), in which a peptide bond was formed between the carboxyl carbon of the AsLOV2 Leu144 and the nitrogen of the amino group of CHT7_CXC's Ala1. The fusions were then docked onto the 12mer TTTGAA containing DNA structure using HADDOCK 2.4 (11, 12) in a similar manner as was done for CHT7_CXC-DNA complex docking (see the section "**Protein and DNA docking"**).

**Generation of $LOV_L$ conformational state**

Photoactivation of AsLOV2 dark state induces the formation of a covalent adduct between the FMN and Cys450. Therefore, to generate the light induced state of AsLOV2 ($LOV_L$) the simulation box was created using the tleap program of the molecular dynamics package Amber (ver20) . The ff14SBonlysc force field (24) was used. Library file and the force field parameter files for the modified FMN (to form covalent adduct) were used as provided by Iuliano et al., 2021 (25). The OPC nonpolarized 4-point 3-charge rigid water model (26) was used. The dark state structure of AsLOV2 (PDBID: 2V1A) and water molecule coordinates were loaded. Then a covalent bond was added between C450(SG) and FMN(C4A). The system was then solvated by adding 5130 water molecules in a truncated octahedral box imposing the restriction that none of the solute molecules approach closer than 8Å to the boundaries of the simulation box.

Further the modeled system was subjected to a minimization run of 100000 steps. During which the whole system (except the H atoms, water molecules, FMN and Cys450) was kept restrained using the force constant of 100 kcal $mol^{-1}$ $Å^{-2}$. Then, using the NVT ensemble, the temperature was increased from 100 K to 298 K within 1 ns (time resolution 1 fs). SHAKE was applied on all the bonds. Particle mesh ewald method (27) was utilized to apply a nonbonded cutoff of 8 Å. Maintaining the restraint, NPT simulation was run for 1ns at 298 K (temperature maintained using Langevin thermostat collision frequency of 1 $ps^{-1}$) and pressure coupling constant was kept as 0.1. Then, after changing the restraint force constant to 10.0 kcal $mol^{-1}$ $Å^{-2}$ and a pressure coupling constant to 0.5, an equilibration run of 1 ns was carried out. Next, the restraints were only applied on the backbone atoms (CA, N, and C) of the protein and the system was minimized for 10,000 steps and then 1 ns NPT simulation was run. During this simulation, all the restraints were maintained as previously, except on C450. This was followed by two run of 1ns after reducing the restraint force constant first to 1.0 kcal $mol^{-1}$ $Å^{-2}$ and then to 0.1 kcal $mol^{-1}$ $Å^{-2}$. Then the system was equilibrated at 298K for another 1 ns, after removing all the restraints. The temporal resolution was reduced to 4 fs by applying hydrogen mass repartitioning (28) and the simulation was extended till 10 μs, which provided us the light induced state of AsLOV2 with completely unfolded $J_\alpha$ helix. All the simulations were carried out using the NVIDIA GPU GTX1660 Ti card.

### Molecular Dynamics simulations of LOV_CXC fusions

DNA docked fusion complexes of $LOV_D$-CXC and $LOV_L$-CXC were simulated using the similar methodology as used for CXC-DNA complexes. Forcefield and parameter files for FMN (in $LOV_D$-CXC) and the modified FMN (in $LOV_L$-CXC) were obtained from the previous work by Iuliano et al., 2021 (25). Whereas, to parameterize CXC and DNA the fusions, ff14SB and bsc1 the force fields were used as mentioned earlier in the methodology for CXC-DNA complex simulations.

### MD simulation analysis

All analyses were carried out using Amber cpptraj (29), Bio3D (30) tool integrated into R (version 4.1.3, (31)) and Pymol (version 2.3.0n (32)) . The simulation trajectories were visualized using VMD (33). Binding energy estimations were made using MMPBSA (34)and changes in DNA groove parameters (such as width, depth, ion concentration) were estimated using the software suite CURVES+ (35, 36).

Refernces:


1. C.-H. Tsai, *et al.*, The protein Compromised Hydrolysis of Triacylglycerols 7 (CHT7) acts as a repressor of cellular quiescence in Chlamydomonas. *Proceedings of the National Academy of Sciences* **111**, 15833–15838 (2014).
2. A. Kanhere, M. Bansal, A novel method for prokaryotic promoter prediction based on DNA stability. *BMC Bioinformatics* **6**, 1 (2005).
3. V. Rangannan, M. Bansal, High-quality annotation of promoter regions for 913 bacterial genomes. *Bioinformatics* **26**, 3043–3050 (2010).
4. A. Sharma, D. T. Nair, MsDpo4-a DinB Homolog from Mycobacterium smegmatis-Is an Error-Prone DNA Polymerase That Can Promote G:T and T:G Mismatches. *J Nucleic Acids* **2012**, 285481 (2012).
5. A. Sharma, J. Kottur, N. Narayanan, D. T. Nair, A strategically located serine residue is critical for the mutator activity of DNA polymerase IV from Escherichia coli. *Nucleic Acids Res* **41**, 5104–5114 (2013).
6. S. Nirwal, D. S. Kulkarni, A. Sharma, D. N. Rao, D. T. Nair, Mechanism of formation of a toroid around DNA by the mismatch sensor protein. *Nucleic Acids Res* **46**, 256–266 (2018).
7. L. M. Hellman, M. G. Fried, Electrophoretic Mobility Shift Assay (EMSA) for Detecting Protein-Nucleic Acid Interactions. *Nat Protoc* **2**, 1849–1861 (2007).
8. A. H. Marceau, *et al.*, Structural basis for LIN54 recognition of CHR elements in cell cycle-regulated promoters. *Nat Commun* **7**, 1–11 (2016).
9. , I-TASSER: a unified platform for automated protein structure and function prediction | Nature Protocols (October 16, 2023).
10. L. P. P. Patro, A. Kumar, N. Kolimi, T. Rathinavelan, 3D-NuS: A Web Server for Automated Modeling and Visualization of Non-Canonical 3-Dimensional Nucleic Acid Structures. *Journal of Molecular Biology* **429**, 2438–2448 (2017).
11. C. Dominguez, R. Boelens, A. M. J. J. Bonvin, HADDOCK: A Protein−Protein Docking

Approach Based on Biochemical or Biophysical Information. *J. Am. Chem. Soc.* **125**, 1731–1737 (2003).
12. R. V. Honorato, *et al.*, Structural Biology in the Clouds: The WeNMR-EOSC Ecosystem. *Frontiers in Molecular Biosciences* **8** (2021).
13. E. F. Pettersen, *et al.*, UCSF Chimera--a visualization system for exploratory research and analysis. *J Comput Chem* **25**, 1605–1612 (2004).
14. D. A. CASE, *et al.*, The Amber Biomolecular Simulation Programs. *J Comput Chem* **26**, 1668–1688 (2005).
15. R. D. Malmstrom, C. T. Lee, A. T. Van Wart, R. E. Amaro, Application of Molecular-Dynamics Based Markov State Models to Functional Proteins. *J Chem Theory Comput* **10**, 2648–2657 (2014).
16. M. K. Scherer, *et al.*, PyEMMA 2: A Software Package for Estimation, Validation, and Analysis of Markov Models. *J Chem Theory Comput* **11**, 5525–5542 (2015).
17. Y. Naritomi, S. Fuchigami, Slow dynamics in protein fluctuations revealed by time-structure based independent component analysis: the case of domain motions. *J Chem Phys* **134**, 065101 (2011).
18. G. Pérez-Hernández, F. Paul, T. Giorgino, G. De Fabritiis, F. Noé, Identification of slow molecular order parameters for Markov model construction. *J Chem Phys* **139**, 015102 (2013).
19. Some Methods for Classification and Analysis of MultiVariate Observations | BibSonomy (July 29, 2023).
20. J.-H. Prinz, *et al.*, Markov models of molecular kinetics: Generation and validation. *The Journal of Chemical Physics* **134**, 174105 (2011).
21. S. Röblitz, M. Weber, Fuzzy spectral clustering by PCCA+: application to Markov state models and data classification. *Adv. Data Anal. Classif.* **7**, 147–179 (2013).
22. Epock: rapid analysis of protein pocket dynamics | Bioinformatics | Oxford Academic (October 17, 2023).
23. A. S. Halavaty, K. Moffat, N- and C-terminal flanking regions modulate light-induced signal transduction in the LOV2 domain of the blue light sensor phototropin 1 from Avena sativa. *Biochemistry* **46**, 14001–14009 (2007).
24. J. A. Maier, *et al.*, ff14SB: Improving the accuracy of protein side chain and backbone parameters from ff99SB. *J Chem Theory Comput* **11**, 3696–3713 (2015).
25. J. N. Iuliano, *et al.*, Unraveling the Mechanism of a LOV Domain Optogenetic Sensor: A Glutamine Lever Induces Unfolding of the Jα Helix. *ACS Chem Biol* **15**, 2752–2765 (2020).
26. S. Izadi, R. Anandakrishnan, A. V. Onufriev, Building Water Models: A Different Approach. *J Phys Chem Lett* **5**, 3863–3871 (2014).
27. T. Darden, D. York, L. Pedersen, Particle mesh Ewald: An N· log(N) method for Ewald sums in large systems. *The Journal of Chemical Physics* **98**, 10089–10092 (1993).
28. C. W. Hopkins, S. Le Grand, R. C. Walker, A. E. Roitberg, Long-Time-Step Molecular Dynamics through Hydrogen Mass Repartitioning. *J Chem Theory Comput* **11**, 1864–1874 (2015).
29. D. R. Roe, T. E. I. Cheatham, PTRAJ and CPPTRAJ: Software for Processing and Analysis of Molecular Dynamics Trajectory Data. *J. Chem. Theory Comput.* **9**, 3084–3095 (2013).
30. B. J. Grant, A. P. C. Rodrigues, K. M. ElSawy, J. A. McCammon, L. S. D. Caves, Bio3d: an R package for the comparative analysis of protein structures. *Bioinformatics* **22**, 2695–2696 (2006).
31. R: a language and environment for statistical computing (July 29, 2023).
32. PyMOL | pymol.org (July 29, 2023).
33. W. Humphrey, A. Dalke, K. Schulten, VMD: visual molecular dynamics. *J Mol Graph* **14**, 33–38, 27–28 (1996).
34. B. R. Miller, *et al.*, MMPBSA.py: An Efficient Program for End-State Free Energy Calculations. *J Chem Theory Comput* **8**, 3314–3321 (2012).
35. C. Blanchet, M. Pasi, K. Zakrzewska, R. Lavery, CURVES+ web server for analyzing and visualizing the helical, backbone and groove parameters of nucleic acid structures. *Nucleic Acids Res* **39**, W68–W73 (2011).

36. R. Lavery, J. H. Maddocks, M. Pasi, K. Zakrzewska, Analyzing ion distributions around DNA. *Nucleic Acids Res* **42**, 8138–8149 (2014).

**Fig.S1.**

A)

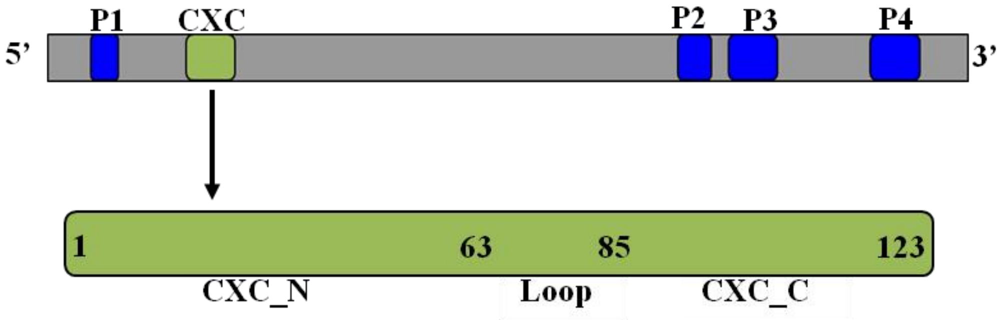


B)

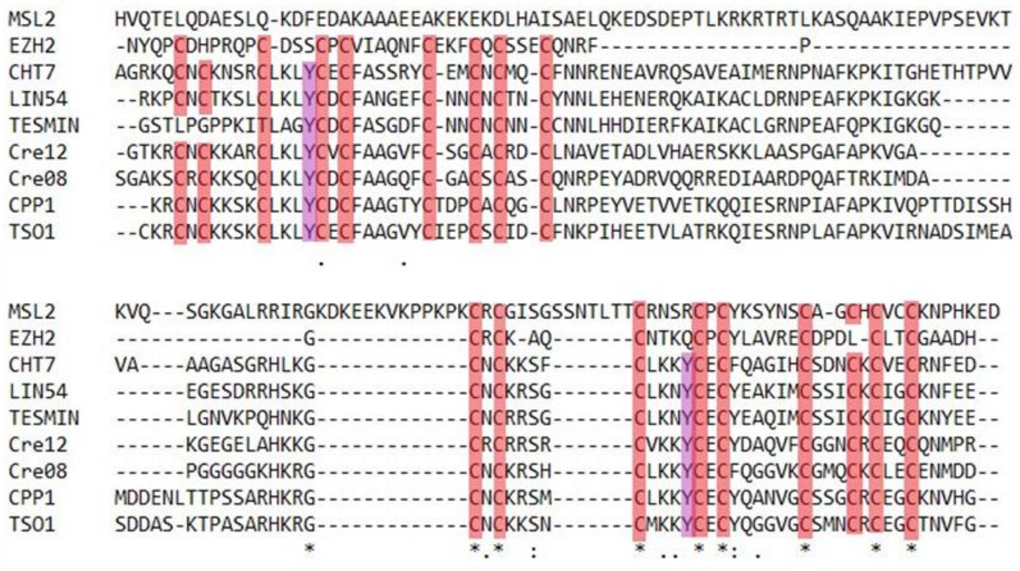


**Fig.S1: A)** Domain arrangement of CHT7. Four predicted protein domains (PPDs: P1, P2, P3, P4) are shown in blue and a DNA binding CXC domain (CHT7_CXC) in green. CXC’s N-terminal (CXC_N), Loop and C-terminal (CXC_C) subdomains further shown as per their residue wise positioning. **B)** Sequence alignment of CXC domains of different proteins. MSL2 from Drosophila melanogaster, EZH2 from humans, CHT7’s (Cre11.g481800.t1.1 ) CXC domain from *Chlmydomonas reinhardtii*, LIN54 from humans, TESMIN from Mouse, Cre12 (Cre12.g550250.t1.2) and Cre08 (Cre08.g361400.t1.2) are other two CXC domain containing proteins in *Chlmydomonas reinhardtii*, CPP1 from Soybean, TSO1 from higher plants. Cysteines within the CXC subdomains are highlighted in red and the subdomain tyrosine’s are in purple. * depict conserved residues.

**Fig.S2.**

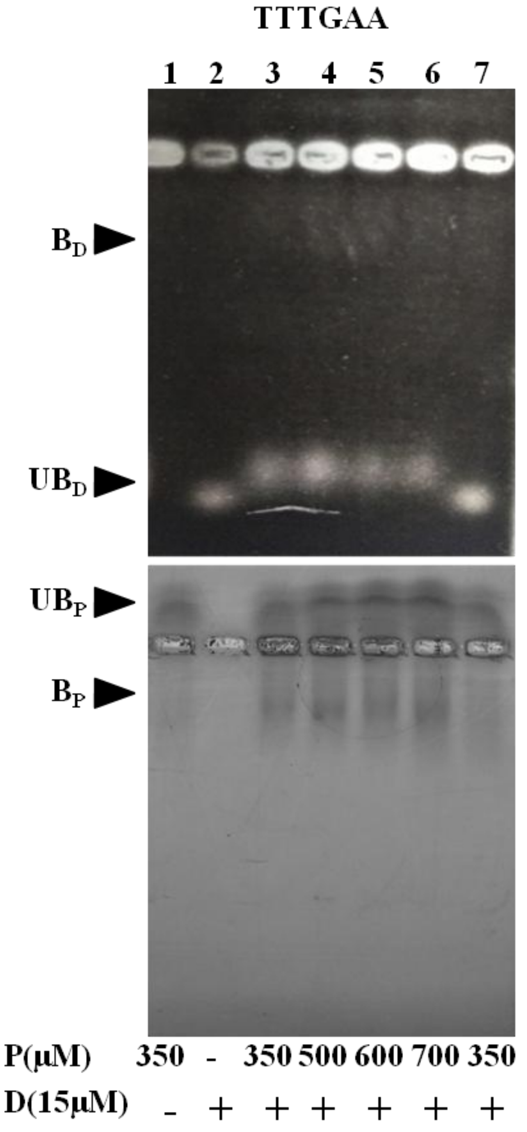


**Fig.S2:** Images of EMSA gel stained for DNA (upper panel) and for protein (lower panel) showing in Lane 1-6 CHT7_CXC weak binding towards 13mer DNA duplex 5'-GGTGTTTGAATTT-3' and in Lane 7 towards the random DNA sequence 5'GATCAATAGATGCGCAGATCATTTACATT-3'. Concentrations of Protein and DNA used are as written in the bottom. Bound and unbound states for DNA (or Protein) are marked as $B_D$ ($B_P$) and $UB_D$ (or $UB_P$), respectively.

**Fig.S3.**

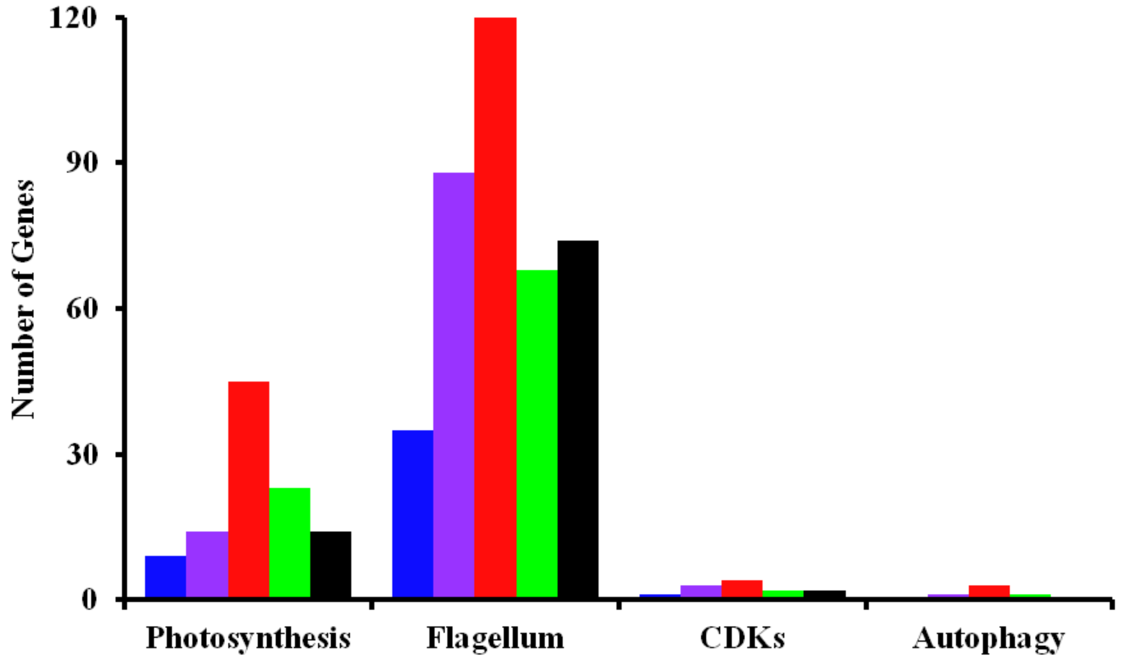


**Fig.S3:** Bar graph shows the number of mis regulated Photosynthetic, Flagellum, CDKs and Autophagy genes in the cht7 mutant that contains six nucleotide (TTYRAA, TTTGAA, CCTGAA, CCTGCC and TTTGCC) sites in the promoter regions within1000 bps upstream of the 5'UTR. Bars for the respective sites are shown in black, blue, purple, red and green. In TTYRAA site, Y depicts Pyrimidines and R depicts Purines.

**Fig.S4.**

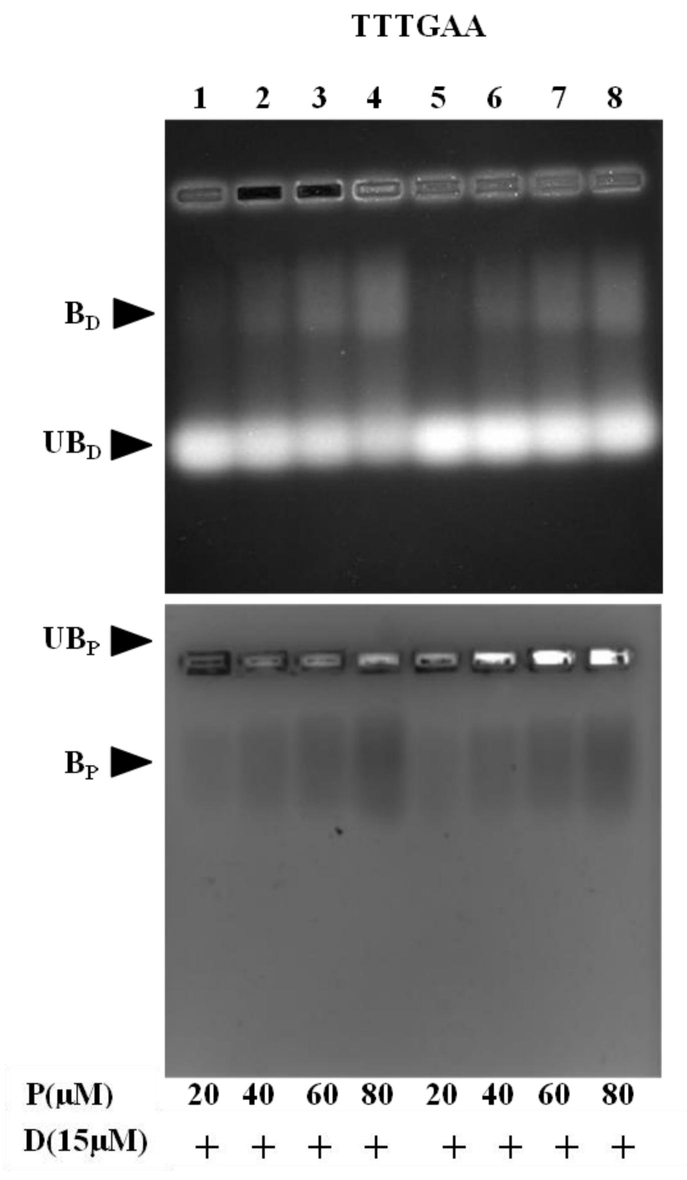


**Fig.S4:** Images of EMSA gel stained for DNA (upper panel) and for protein (lower panel) showing binding of CHT7_CXC to the TTTGAA site within the mis regulated photosynthetic genes, namely LHCA6 (5'-TGGACTGGATTTGAACGATTGAAAGCT-3', Lane 1-4) and LHCB5 (5'-TTCTTTTCATTTGAATGGTTTTCCGAC-3', Lane 5-8). Concentrations of Protein and DNA used are as written in the bottom. Bound and unbound states for DNA (or Protein) are marked as $B_D$ ($B_P$) and $UB_D$ (or $UB_P$), respectively.

**Fig.S5.**

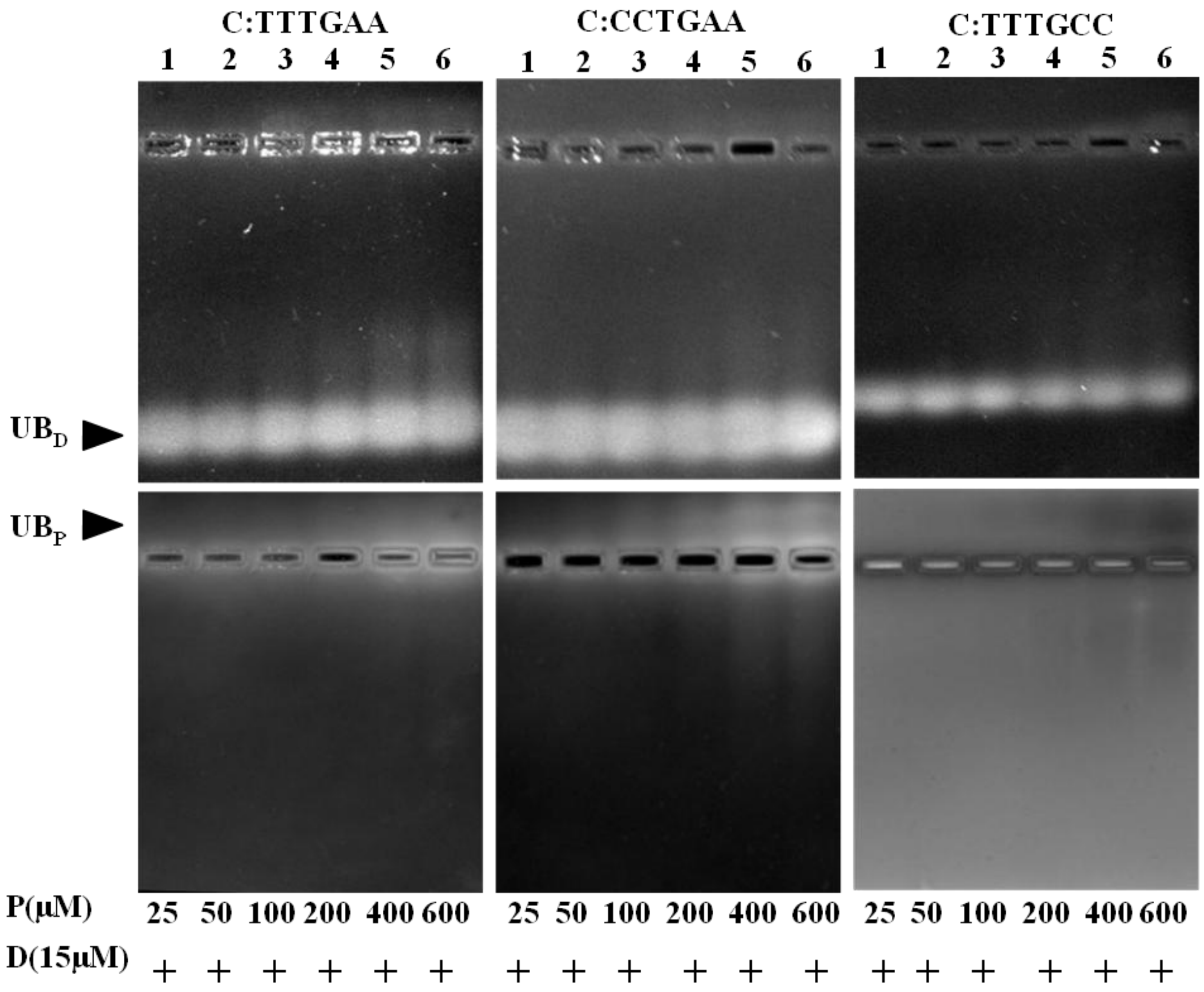


**Fig.S5:** Images of EMSA gels stained for DNA (upper panel) and for protein (lower panel) showing CXC_C binding reactions for 27mer DNA duplex having A) TTTGAA, B) CCTGAA and C) TTTGCC as the binding regions. Concentrations of Protein and DNA used are as written in the bottom. Only unbound states of the DNA and the protein are visible and are marked with UB$_D$ and UB$_P$, respectively.

**Fig.S6.**

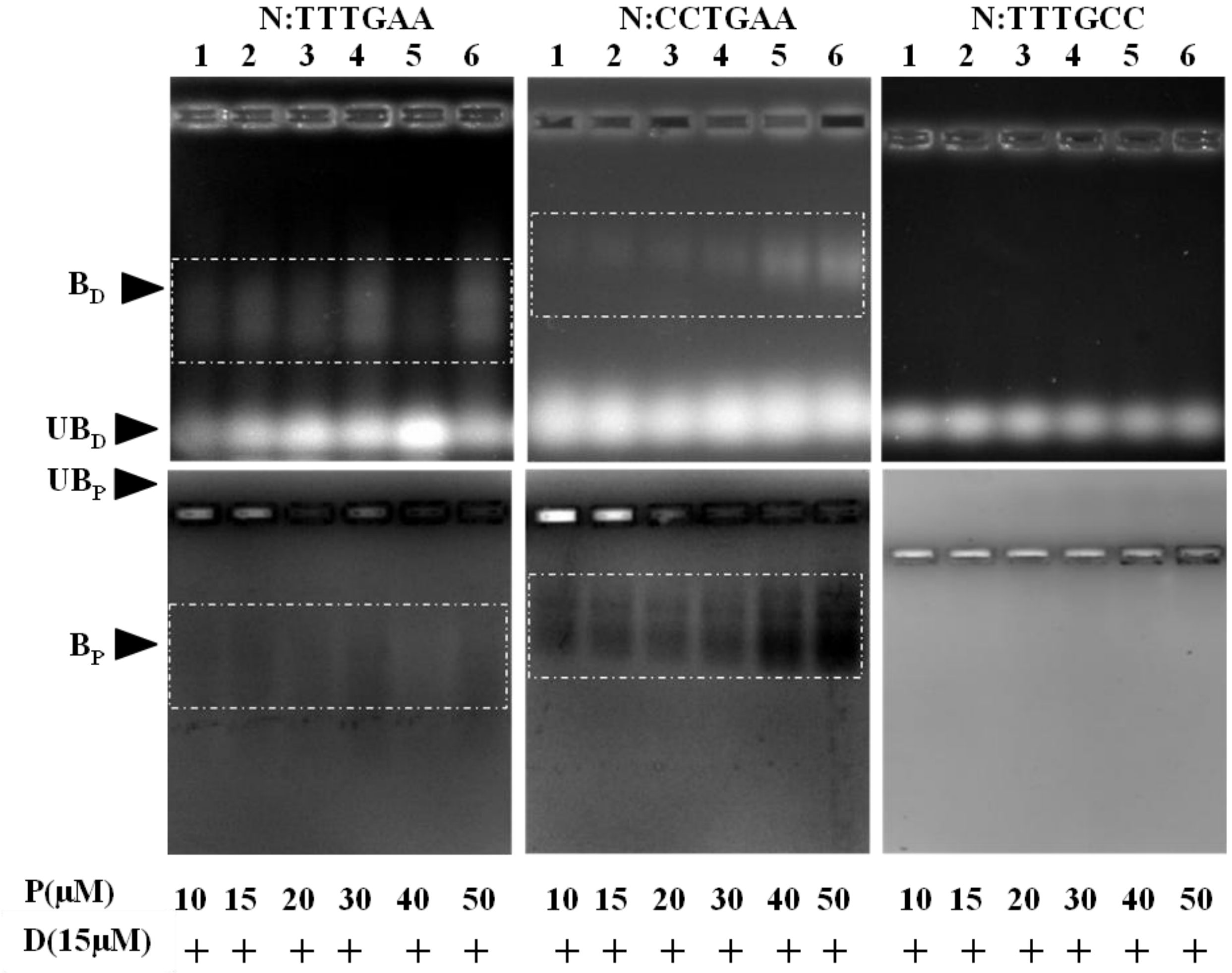


**Fig.S6:** Images of EMSA gels stained for DNA (upper panel) and for protein (lower panel) showing CXC_N binding reactions for 27mer DNA duplex having A) TTTGAA, B) CCTGAA and C) TTTGCC as the binding regions. Bound states seen in case A) and B) are marked in dotted rectangles. Concentrations of Protein and DNA used are as written in the bottom. Bound and unbound states for DNA (or Protein) are marked as $B_D$ ($B_P$) and $UB_D$ (or $UB_P$), respectively.

**Fig.S7.**

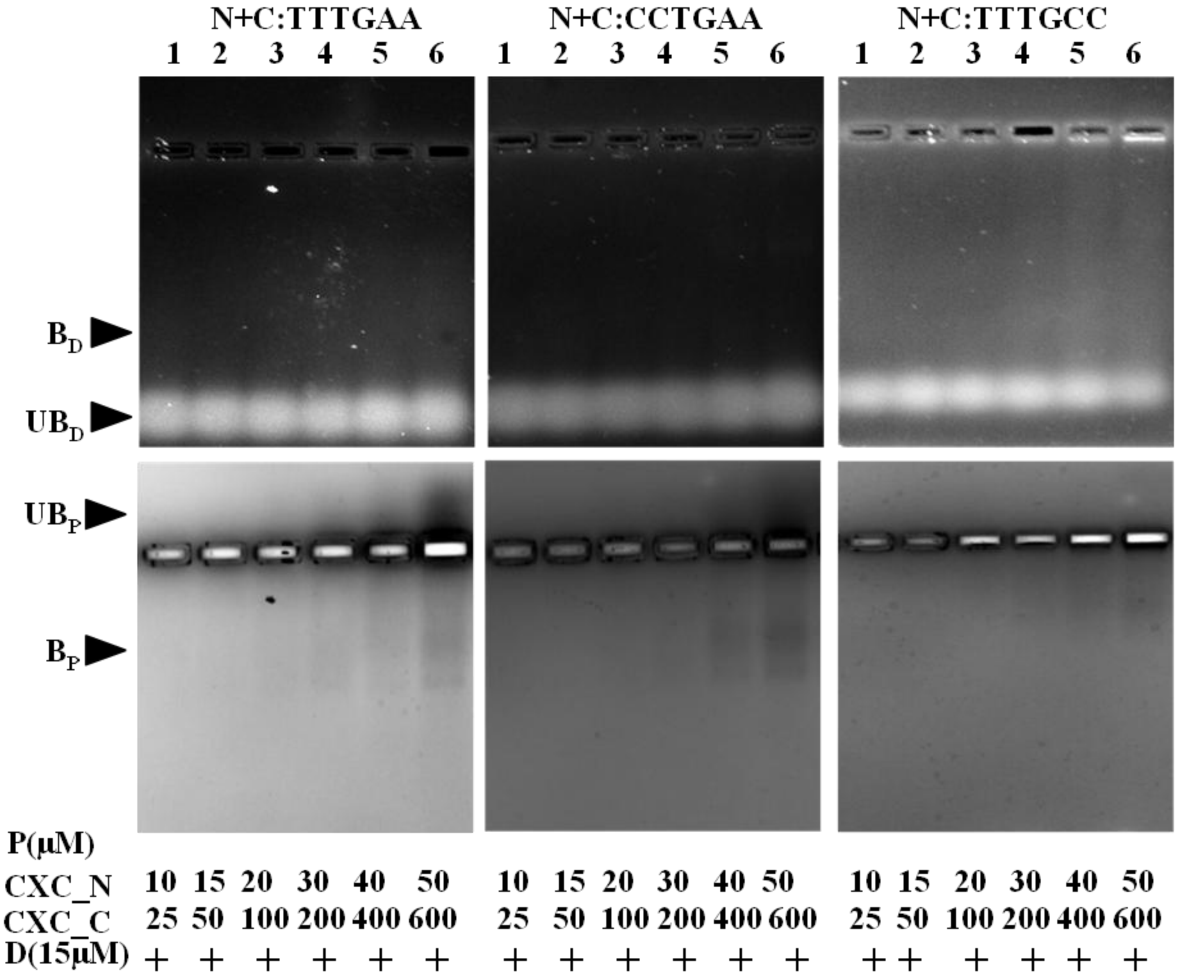


**Fig.S7:** Images of EMSA gels stained for DNA (upper panel) and for protein (lower panel) showing binding reactions when both the subdomain proteins (CXC_N and CXC_C) were taken together to bind onto the 27mer DNA duplex having A) TTTGAA, B) CCTGAA and C) TTTGCC as the DNA binding regions. Smeared signal for DNA and protein is marked by $B_D$ and $B_P$ and is indicative of the presence of weakly bound protein DNA complexes that separate during the EMSA run. Concentrations for both the subdomains and DNA used are as written at the bottom. Unbound states of DNA and protein are marked by $UB_D$ and $UB_P$, respectively.

**Fig.S8.**

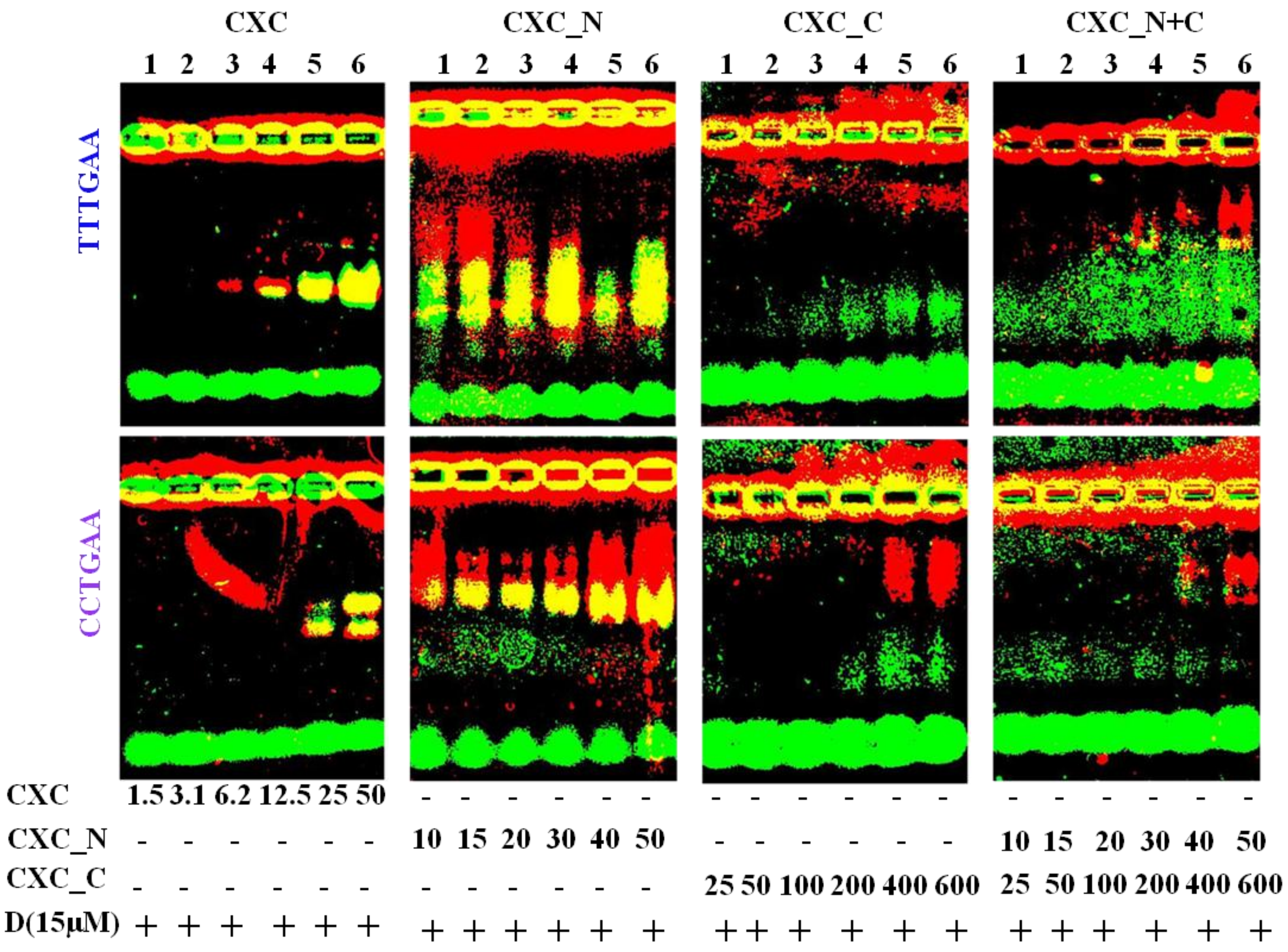


**Fig.S8:** Overlap of EMSA gel images stained for DNA (in green) and for protein (in red). Regions in yellow represent the protein DNA bound complex. Each image shows the protein (CXC, CXC_N, CXC_C and CXC_N+C (CXC_N + CXC_C)) titration against 27mer DNA duplexes containing TTTGAA (upper panel) and CCTGAA (lower panel) as DNA binding sites. Concentration of protein and DNA used is written at the bottom.

**Fig.S9.**

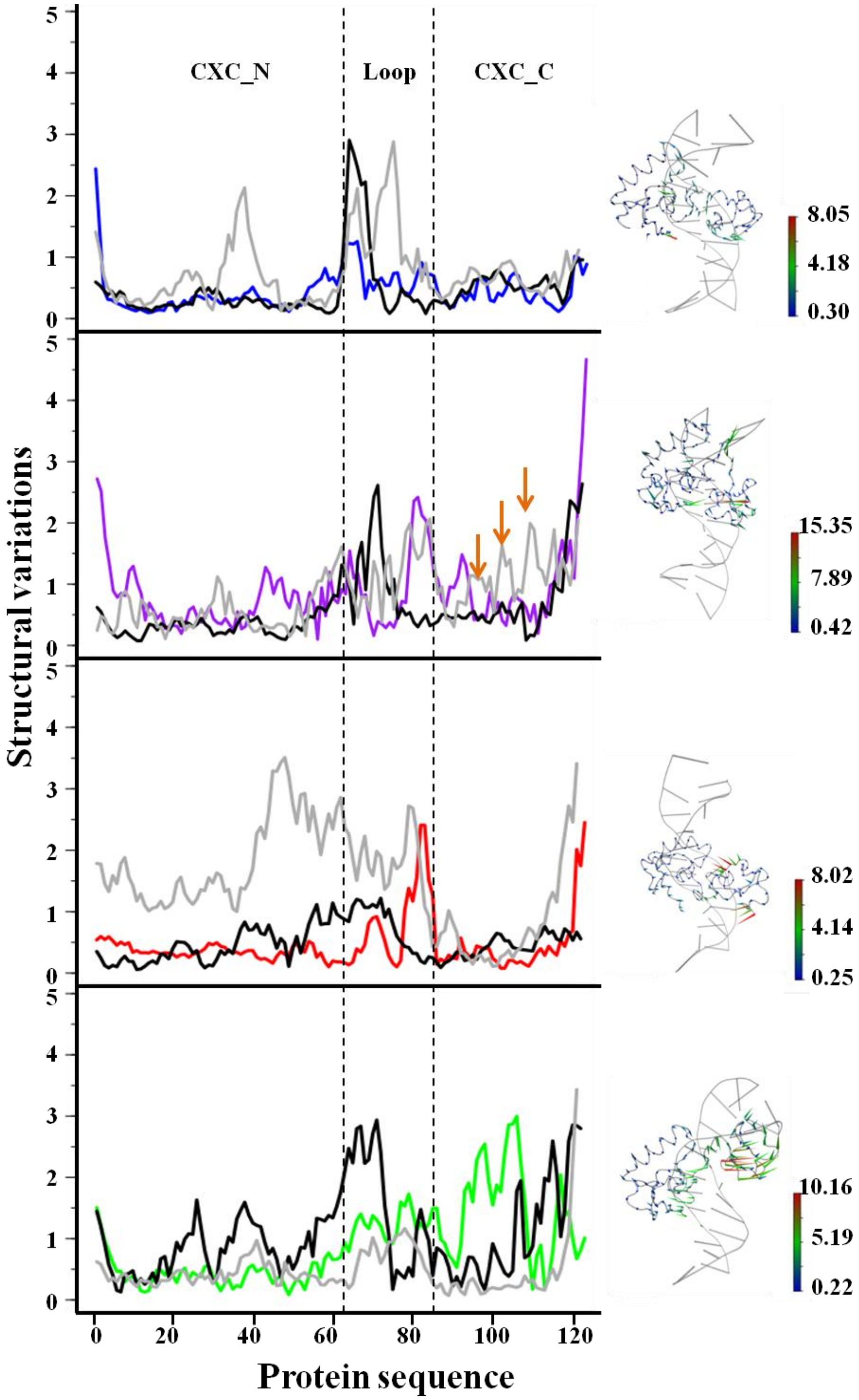


**Fig.S9:** Residue wise structural variations within the first principal component (pc1) of the CHT7_CXC structure in complex with DNA containing binding regions TTTGAA, CCTTGAA, CCTGCC and TTTGCC are plotted in blue, purple, red and green, respectively. Also, shown (on right) are the porcupine models for these complexes. Where, the scale bar shows the extent of the structural variations. Variations in each complex are compared with the variations due to the corresponding complexes formed using LIN54 (black) and LIN54 whose loop domain is swapped with that of CHT7_CXC (grey). Increase in structural variations in LIN54 due to the swapped loop is marked by orange arrows in case of CCTGAA plots.

**Fig.S10.**

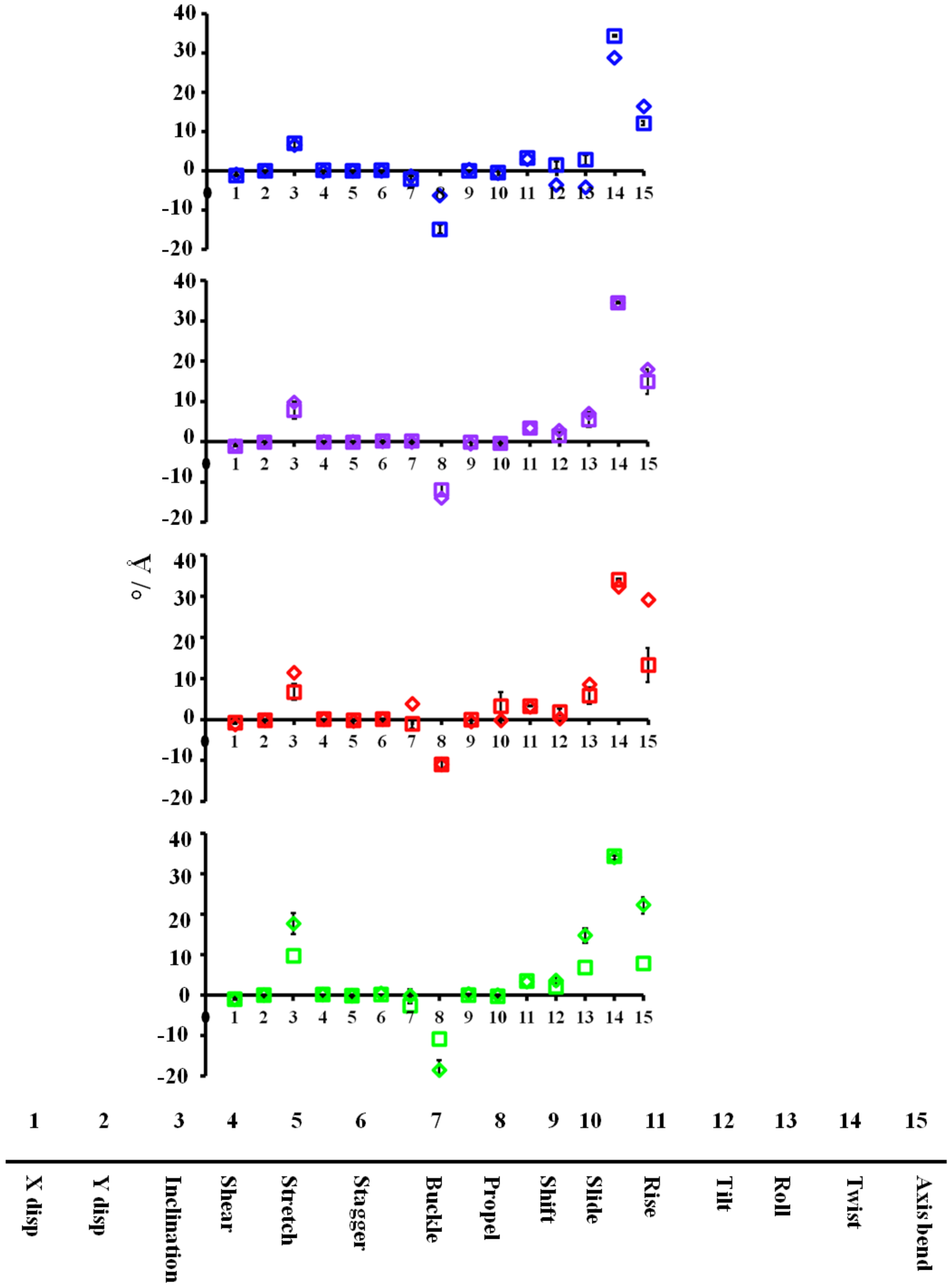


**Fig.S10:** Plotted are the mean values of 15 different DNA base pair geometric parameters estimated from the bound CHT7_CXC complexes (open square) with the 27mer DNA sequences containing TTTGAA (blue), CCTGAA (purple), CCTGCC (red) and TTTGCC (green) as the binding sites. Comparison is shown with the values estimated from the isolated DNA sequences (open rhombus). Bars are the standard errors of the mean estimated from a set of three simulations of CHT7_CXC DNA complexes.

**Fig.S11.**

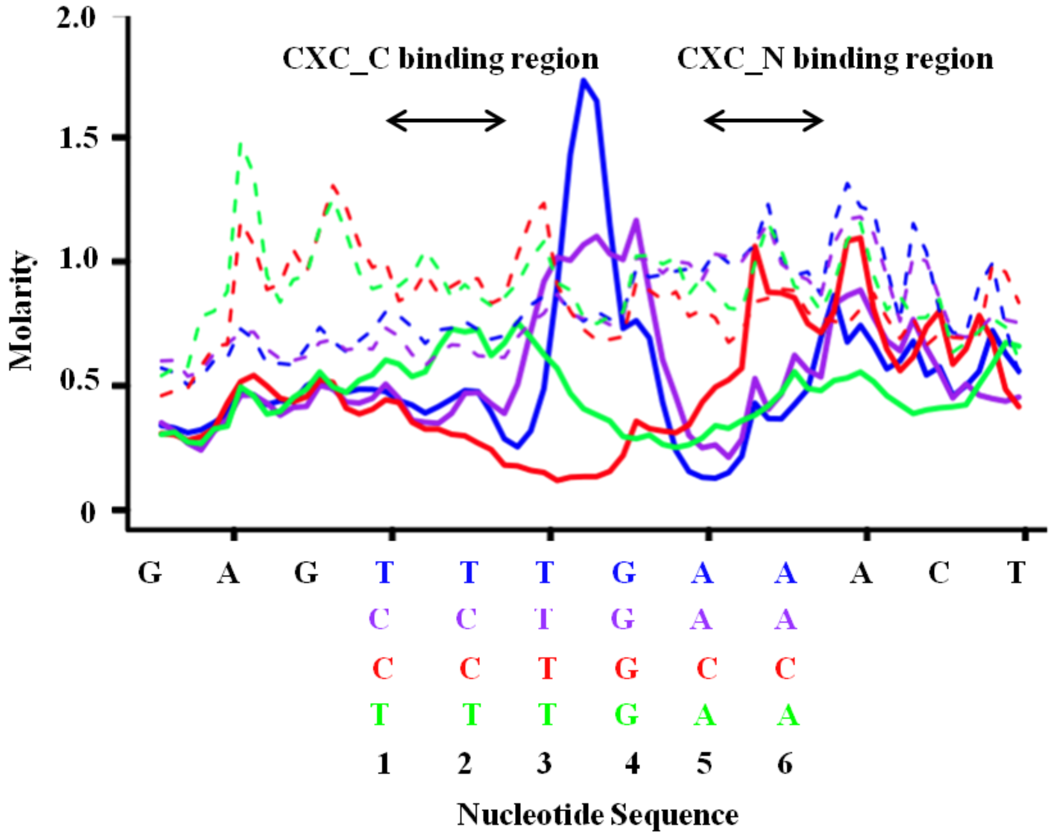


**Fig.S11:** Ionic concentration across the DNA within the CHT7_CXC bound complexes (solid lines) and in isolated DNA (dotted). DNA containing TTTGAA, CCTGAA, CCTGCC and TTTGCC are color coded as blue, purple, red and green, respectively. Sharp dip in ionic concentration in case of TTTGAA and CCTGAA surrounding T1T2 and A5A6 is marked by orange arrows. Increase in ionic concentration at T3G4 positions in TTTGAA and CCTGAA complex is marked by arrows in grey. Subdomain regions that bind at T1T2 and A5A6 nucleotide regions is marked as CXC_C and CXC_N.

**Fig.S12.**

A)

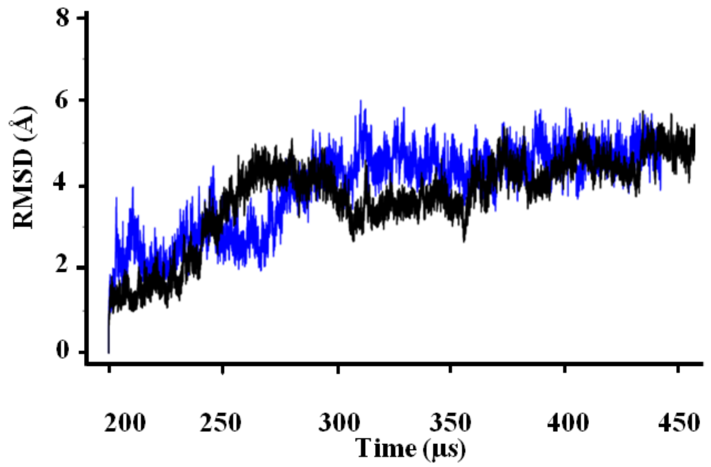


B)

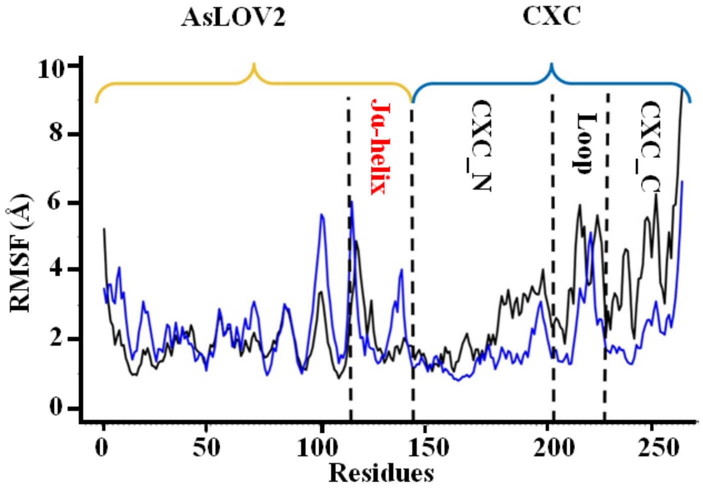


**Fig.S12:** A) RMSD and B) RMSF estimated from the MD simulation data collected over 500 ns for the molecular fusion of *Avena Sativa's* LOV2 (AsLOV2) and CHT7_CXC under the dark ($LOV_D$-CXC) and the light ($LOV_L$-CXC) conditions in complex with 12mer DNA duplex containing TTTGAA as the binding region. Plots for $LOV_D$-CXC and $LOV_L$-CXC are in black and blue, respectively. Regions corresponding to AsLOV2, its $J_\alpha$ helix and CHT7_CXC subdomains and its loop are mentioned in B).

**Fig.S13.**

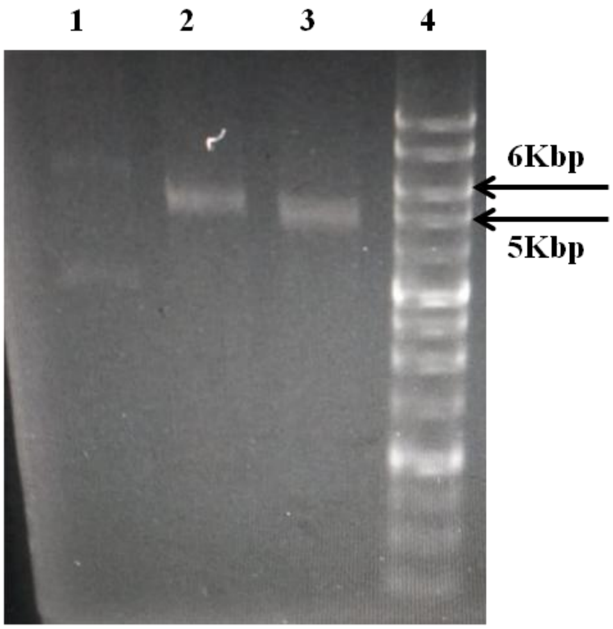


**Fig.S13.** Enzyme digestion of plasmid (Pgex6p1 (5 Kbps) + CHT7_CXC gene insert (384 bps)). Lanes 1: intact plasmid, 2: BamHI digested plasmid, 3: BamHI and XhoI digested plasmid, 4: DNA marker ladder.

**Fig.S14.**

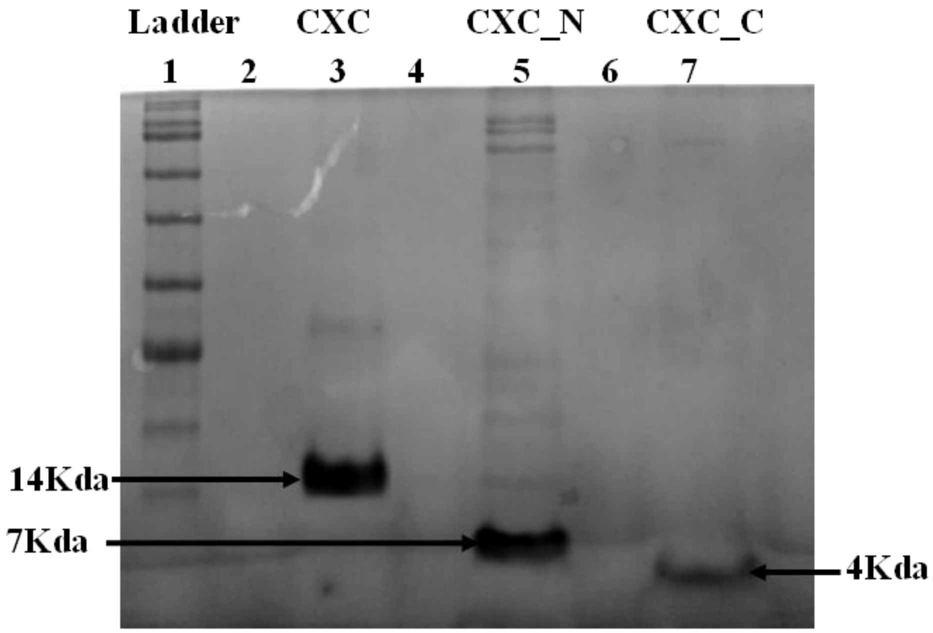


**Fig.S14:** SDS PAGE showing GST column purified CHT7_CXC (14 kDa, lane 3), CXC_N (7 kDa, lane 5), CXC_C (4 kDa, lane 7). Protein marker ladder is in lane 1.

Fig.S15.

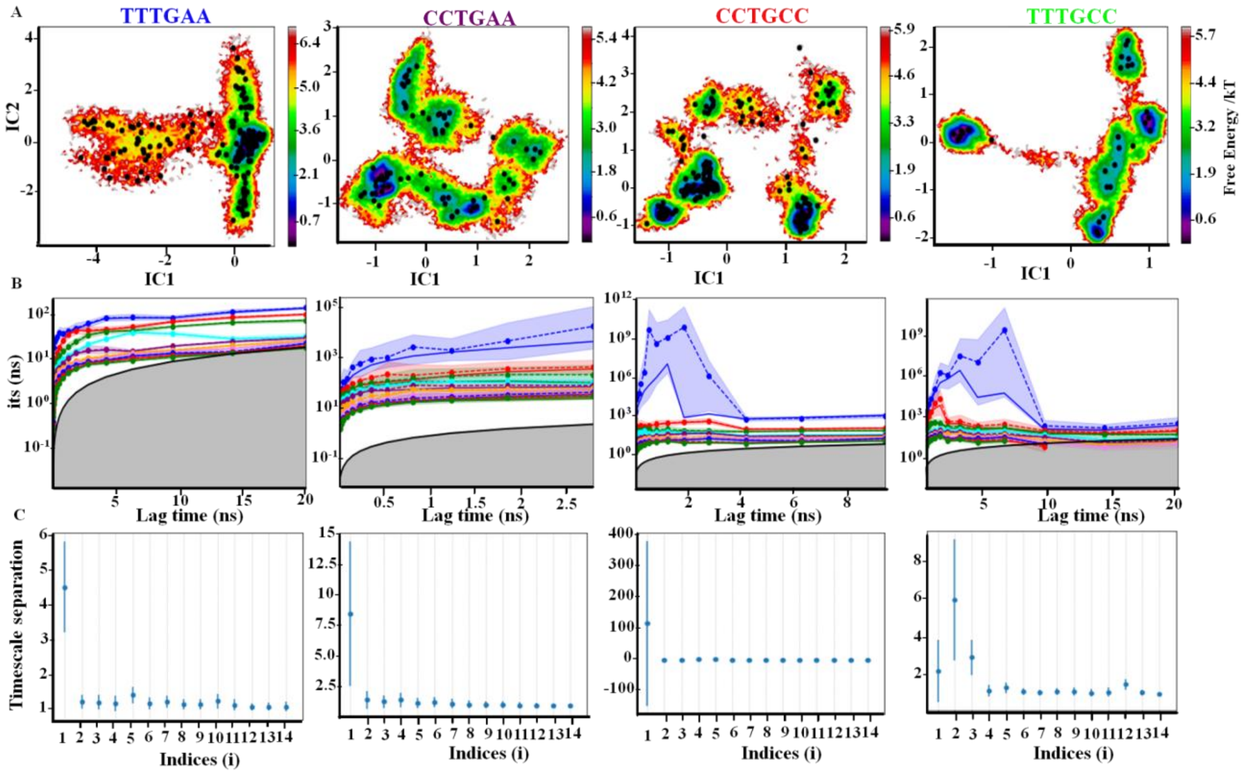


**Fig.S15:** Markov State Modelling associated parameters. **A)** k-mean clusters distributed (black dots) over the free energy map between the two slowest TICA components (IC1, IC2), **B)** Implied relaxation time scales (its) of CHT7_CXC DNA complexes using their respective cluster numbers and **C)** time scale separation ($t_{i+1}/t_i$, where i is index of implied time scale), for TTTGAA, CCTGAA, CCTGCC, TTTGCC complexes, respectively.
